# Identification of Small Molecule Modulators of Diguanylate Cyclase by FRET-based High-Throughput-Screening

**DOI:** 10.1101/402909

**Authors:** Matthias Christen, Cassandra Kamischke, Hemantha D. Kulasekara, Kathleen C. Olivas, Bridget R. Kulasekara, Beat Christen, Toni Kline, Samuel I. Miller

## Abstract

The bacterial second messenger cyclic diguanosine monophosphate (c-di-GMP) is a key regulator of cellular motility, the cell cycle, and biofilm formation with its resultant antibiotic tolerance, which may make chronic infections difficult to treat. Therefore, diguanylate cyclases, which regulate the spatiotemporal production of c-di-GMP, may be attractive drug targets to control biofilm formation that is part of chronic infections. In this paper, we present a FRET-based biochemical high-throughput screening approach coupled with detailed structure-activity studies to identify synthetic small molecule modulators of the diguanylate cyclase, DgcA, from *Caulobacter crescentus*. We identified a set of 7 small molecules that in the low µM range regulate DgcA enzymatic activity. Subsequent structure activity studies on selected scaffolds revealed a remarkable diversity of modulatory behaviors, including slight chemical substitutions that revert the effects from allosteric enzyme inhibition to activation. The compounds identified represent novel chemotypes and are potentially developable into chemical genetic tools for the dissection of c-di-GMP signaling networks and alteration of c-di-GMP associated phenotypes. In sum, our studies underline the importance for detailed mechanism of action studies for inhibitors of c-di-GMP signaling and demonstrate the complex interplay between synthetic small molecules and the regulatory mechanisms that control the activity of diguanylate cyclases.

## Introduction

The second messenger cyclic dimeric guanosine monophosphate (c-di-GMP) mediates diverse bacterial cellular processes including; antibiotic resistance, biofilm formation, extracellular carbohydrate and adhesin production, pilus- and flagellum-based motility, and cell cycle progression (1–5). Signal integration into c-di-GMP networks is, in part, controlled by diguanylate cyclases (DGCs) that convert two molecules of GTP to c-di-GMP. The enzymatic activity of DGCs resides within a conserved domain comprised of the amino acids, glycine-glycine-aspartate-glutamate-phenylalanine (GGDEF) that forms the enzymatic active site. The GGDEF domain shares similarity to the PALM 4 domain found in other classes of nucleotide cyclase (6–10). The ap-parent role of c-di-GMP in the cell cycle and the presence of many paralogous DGC enzymes controlling diverse cellular functions indicate that there is likely tight spatial and temporal regulation of c-di-GMP (10–12). Bacterial genomes encode multiple GGDEF domains in proteins with signal-sensing domains (13). However, the presence of multiple paralogs makes it difficult to study the signaling processing properties of c-di-GMP signaling networks using conventional genetic techniques. Therefore, chemical genetic approaches to inactivate each segment of the c-di-GMP signaling network may be an attractive approach to study the overall biological function of c-di-GMP. In pathogenic bacteria, cellular production of c-di-GMP is essential to maintain biofilm formation, especially under stressed conditions such as induction by aminoglycoside antibiotics (1). Small molecules that effectively inhibit DGC activity have the potential to prevent biofilm formation, thus, making DGCs interesting targets to develop new classes of antimicrobial agents.

We developed a sensitive and robust FRET based DGC activity assay and performed high-throughput (HT) screening on a comprehensive compound library to evaluate 27,502 small molecules for inhibition of the *Caulobacter crescentus* DGC DgcA (CC3285). DgcA has been extensively characterized and therefore serves as a model enzyme to study c-di-GMP related phenotypic effects (9, 11, 12, 14). Similar to the majority of DGC enzymes, DgcA is subjected to high affinity binding of c-di-GMP to an allosteric site (I-site), which efficiently blocks enzymatic activity in a non-competitive manner and resides distant from the catalytic pocket (A-site). Mutational analysis of DgcA has provided convincing evidence that c-di-GMP binding to several conserved charged amino acids at the I-site is a key mechanism for allosteric regulation of DGCs (9).

## Results

### Development of a FRET-based biochemical high-throughput screen to monitor c-di-GMP production

To identify small molecule inhibitors of DgcA, we established a sensitive FRET-based activity assay. We previously reported a fluorescence resonance energy transfer (FRET)–based c-di-GMP biosensor, which we refer to herein as biosensor (10). This biosensor consists the c-di-GMP binding domain of the PilZ protein YcgR, which is C-and Nterminally tagged with mCYPet and mYPet fluorescent proteins. Binding of c-di-GMP to the YcgR domain induces a conformational change that alters the relative orientation of the external fluorescent subunit protein reporters leading to a decrease in FRET efficiency. Thus, the fluorescence properties (535/470nm emission ratio) of the biosensor reflect c-di-GMP levels. The biosensor exhibits a change in net FRET (nFRET) of –60.6%, with a binding constant of 198 nM for c-di-GMP and no detectable response to cyclic adenosine 3,5-monophosphate, cyclic GMP, or guanosine 5-triphosphate (10).

To measure suitability and performance of the biosensor for HT-capable DGC assay, we recorded kinetics of fluorescence emission ratio change in 384-well plates in a 20 µL reaction volume with 20 nM DgcA enzyme and 50 nM biosensor and followed the change in fluorescence in intervals of 1 min (Fig. 1A). Addition of 50 nM DgcA in absence of GTP does not alter FRET efficiency (Fig 1A, open circles) while injection of 20 µM GTP induces a rapid change in FRET efficiency (Fig 1A, closed circles). Based on the FRET ratio for the free and c-di-GMP saturated biosensor, we determined the corresponding increase in c-di-GMP (Fig. 1B). Our sensitive and robust FRET assay detects c-di-GMP production in the nM range and permits determination of initial rate kinetics at levels below occurrence of allosteric c-di-GMP inhibition.

**Fig 1.**
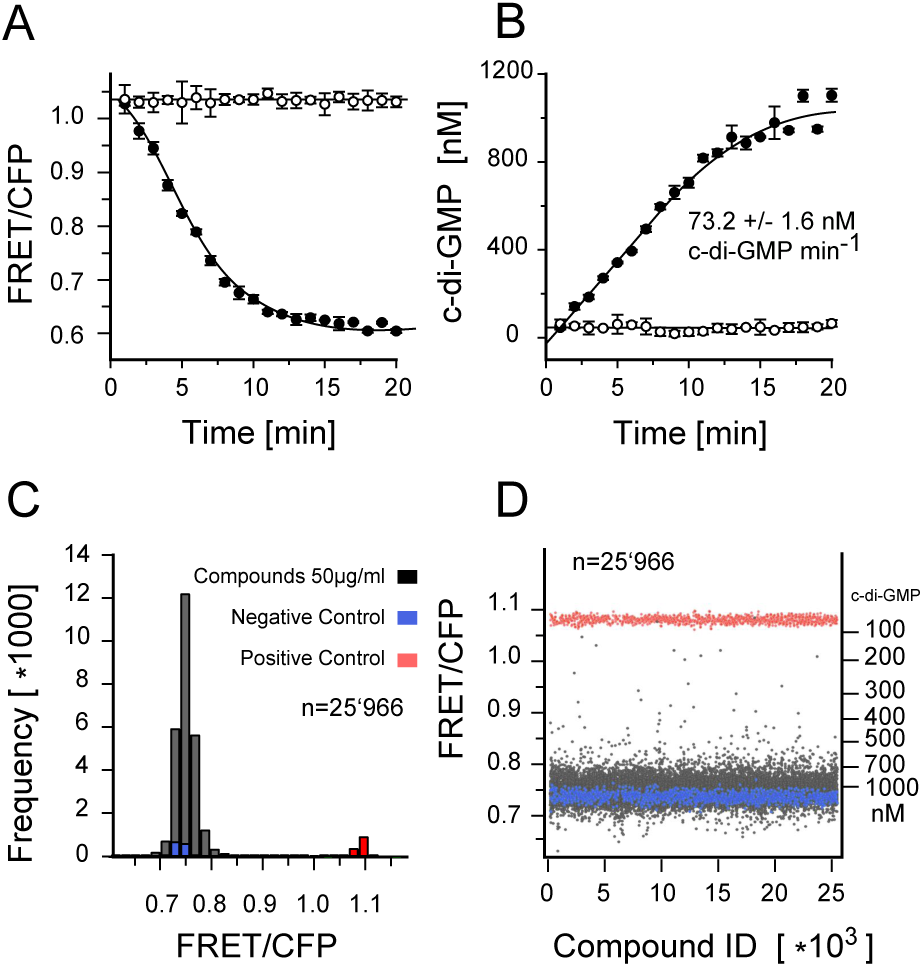
FRET assay for c-di-GMP. (**A**) Kinetics of fluorescence emission ration change (527/480nm) measured in a 384 well plate format. Affinity purified YcgR FRET biosensor was added in presence of 20 nM DgcA enzyme and 20µM GTP substrate (closed circles) or in absence of GTP (open circles). (**B**) Corresponding increase in c-di-GMP concentration derived from the change in fluorescence emission ration (535/470nm). Above 800 nM c-di-GMP, allosteric product inhibition decreases DGC activity of DgcA. Each graph shows the average of three independent measurements. The reaction volume per well was 20 µl in a 384 well Corning low volume flat bottom plate. (**C**) Histogram of the fluorescence emission ratio after 3 hours incubation of 20 nM DgcA with 20 µM GTP in presence of 50µg/ml compounds and 1% DMSO. Wells without GTP substrate added were used as positive controls (red), wells with exogenous c-di-GMP (5 µM) were used as negative controls (blue). (**D**) Plate uniformity of the HT screening. For every well, background fluorescence signal of the compound has been measured prior addition of the FRET-biosensor. Compounds with auto fluorescence exceeding 7% of the FRET biosensor signal (535/470nm) were discarded.

### FRET-based HT-screening and discovery of primary hits for diguanylate cyclase inhibition

A major challenge of biochemical HT screening is to gain significance on large-scale experimental datasets for quality control and hit selection. Through prior screening, we determined plate uniformity, signal variability and repeatability assessment assays for CV and Z-factor calculation. The average Z-factor for 3 plates read on two consecutive days was 0.71, and the CV for high, medium, and low signal were well below 5%. The chemical library screened encompasses 27,505 commercially available compounds derived from small-molecule chemical libraries maintained at the Institute of Chemical Biology (ICCB) at Harvard University, Boston, MA, funded by the Regional Centers of Excellence in BioDefense and Emerging Infectious Disease (NSRB/NERCE). During screening, we first preincubated the DgcA enzyme with 200 nL compound (5 mg/mL) in DMSO and measured fluorescence emission ratio before and after addition of the biosensor to monitor fluorescence of chemical compounds and FRET at 470 and 535 nm. This second measurement will also detect compounds that mimic c-di-GMP agonists. After addition of 20 µM GTP, FRET was measured 3 h after incubation as an endpoint measurement (Fig. 1C,D)) (materials and methods). For a compound to be considered a hit, the following criteria had to be met: (i) Compounds with c-di-GMP production below 300 nM after 3h incubation with DgcA and 20 µM GTP were considered strong hits; (ii) duplicates were consistent with a difference in final c-di-GMP levels <100 nM; and background fluorescence prior to addition of the FRET biosensor must not exceed 7% of the FRET signal in CFP and YFP emission channels. We used total fluorescence as a quality control criterion to identify inherently fluorescent molecules. From the 27,502 compounds screened in duplicates, 1029 (3.7%) were removed from the analysis due to autofluorescence. We identified 49 compounds (0.18%) as strong hits with c-di-GMP levels below 300 nM after 3h incubation with DgcA. An additional 241 compounds (0.87%) were classified as moderate inhibitors with c-di-GMP levels below 400 nM after incubation with DgcA.

### Selection of compound scaffolds for secondary assays and SAR studies

In selecting compounds for structure-activity relationship (SAR) studies, we placed a priority on compounds likely to be functioning via our target mechanism of action. We rejected cytotoxic or cytostatic compounds and favored those compounds that exhibited synthetic tractability and suitability for chemotype expansion. Based on these criteria, we selected 18 compounds out of 49 initial hits for confirmatory and secondary assays. We found that 38.9% (7 out of 18) had an IC_50_ smaller than 50 µM, while 22.2% percent (4 out of 18) also had IC_50_ smaller than 10 µM (Fig. 2A). The results from our secondary assays led us to focus further structure activity studies on the two scaffolds [2-oxo-2-(2-oxopyrrolidin-1-yl)ethyl] 1,3-benzothiazole-6-carboxylate **1** and 4-(2,5-dimethylphenoxy)-N-(4-morpholin-4-ylphenyl)butanamide **2** with IC_50_ of 4.0 (Fig 2B) and 6.4 µM (Fig 2C). The concentration response curve of compound **1** revealed complete inhibition of DgcA activity (Fig. 2B). In contrast, compound **2** acts as partial inhibitor for DgcA (Fig 2C) with remaining 31.5% DGC activity in the presence of an enzyme saturating inhibitor concentration. This indicates that upon binding of **2**, the DgcA enzyme is converted into a modified but still partially functional enzyme–substrate-inhibitor complex raising the possibility that **2** binds to a site distinct from the substrate binding site. For subsequent structure activity studies, we chemically synthesized a set of 24 derivatives of compound **1** and 16 compounds with structural similarities to scaffold **2** (material and methods).

**Fig 2.**
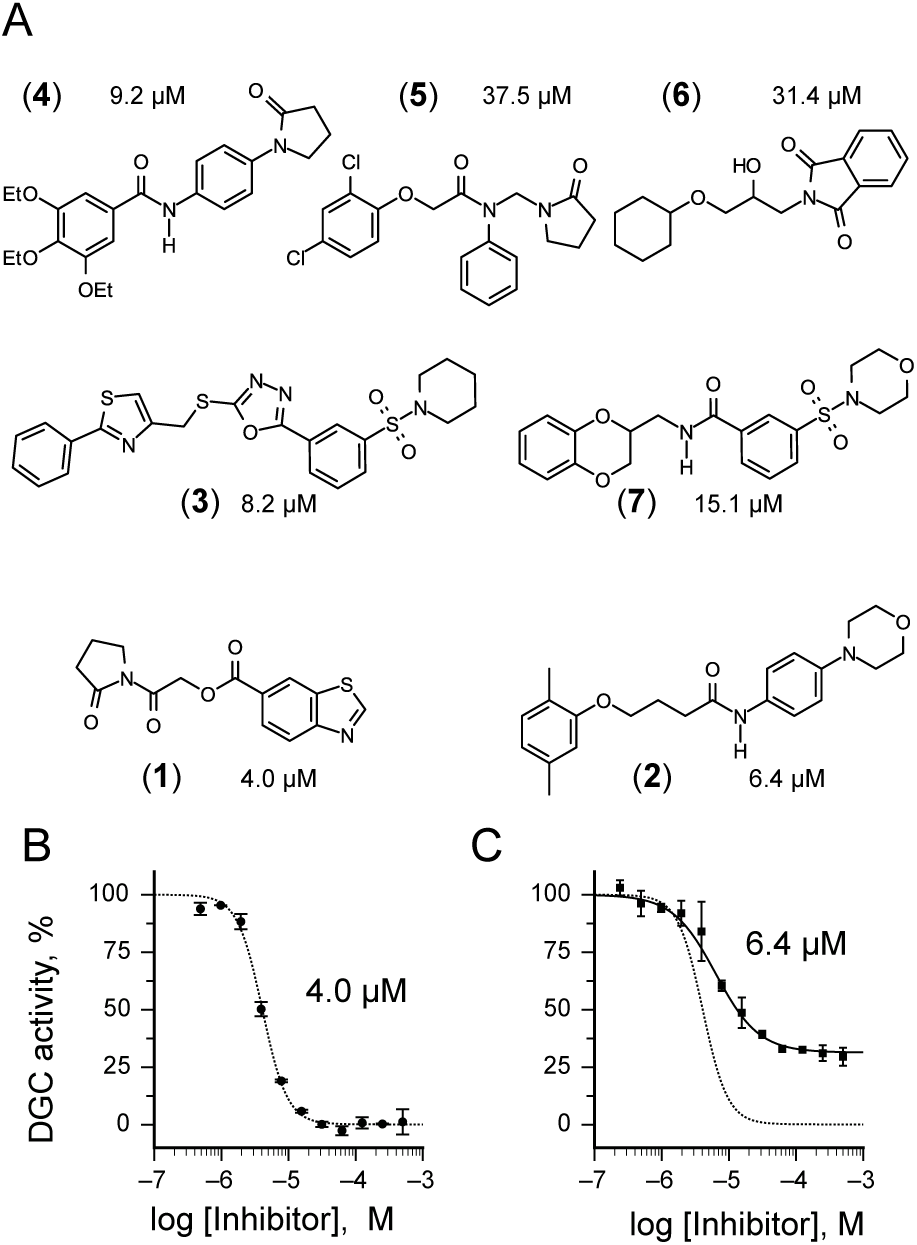
Inhibitors of the DGC enzyme DgcA. (**A**) Structure of the small molecular inhibitors **1**, **2**, **3**, **4**, **5**, **6** and **7** identified in the FRET-screening with IC_50_ values below 50 µM. Out of them, 4 compounds **1**, **2**, **3** and **4** exhibit IC_50_ values <10 µM. Scaffolds **1** and **2** have been selected for subsequent SAR studies. (**B**) The concentration response curve of compound **1** (IC_50_ 4 µM) reveals complete inhibition against DgcA. (**C**) Compound **2** (IC_50_ 6.4 µM) acts as partial inhibitor for DgcA. The corresponding concentration response curve of **1** is shown as dashed line.

### Characterization of structure activity relationship of scaffold 1

Structure activity studies (SAR) can be used to probe steric constraints imposed in the binding site and reveal the molecular relationship between biological activity and structural properties of a ligand. Scaffold **1** contains a 1-acetylpyrrolidin-2-one linked to 1,3-benzothiazole-6-carboxylate (Fig. 2A). We individually probed the pyrrolidin-2-one ring, the linker region and the nucleobase-like 1,3-benzothiazole-6-carboxylate moiety for steric demand and biological activities. Therefore, we tested binding affinities of **1** analogues by replacing the pyrrolidin-2-one ring from R1 and holding the 2-oxoethyl linker and 1,3-benzothiazole-6-carboxylate constant. Our selection of derivatives emphasized pyrrolidine **1a**, dimethylpropanamide **1b**, cyclohexanecarboxamide **1c**, benzamide **1d**, 4-methylaniline **1e** and 2-methoxyaniline **1f** replacing the pyrrolidin-2-one ring. We selected these residues for their steric demand and to probe for hydrogen-bonding, hydrophobic interaction and pi stacking contacts. The inhibitory potencies of the six pyrrolidin-2-one ring replacement analogs are shown in Tab. 1. Many of the derivatives retained but did not exceed a level of potency comparable to that of **1**. For example, replacement of pyrrolidin-2-one with pyrrolidine in **1a** had only marginal effect on binding affinity with a slight increase in IC_50_ from 4 to 17.1 µM (Tab. 1). These results suggest that presence of an H-bond acceptor at the pyrrolidin-2-one in **1** is not a prerequisite for mediating binding affinity to DgcA. Substitution of the pyrrolidin-2-one with dimethylpropanamide in **1b**, does not affect binding affinity (IC_50_ of 3.6 versus 4.0 µM). However, introduction of substituents with increased steric demand such as cyclohexanecarboxamide **1c** or benzamide **1d** show a 10-fold reduction in binding affinity to 47.2 and 48.5 µM respectively. To test whether the decrease in binding affinity is caused by steric hindrance at the binding site, we reduced the steric load and introduced 4-methylaniline **1e** and 2-methoxyaniline **1f**. As expected, we observed an increased in affinity to 8.5 and 10.7 µM respectively. Thus, it seems that the pyrrolidin-2-one ring of scaffold **1** can be replaced by alternative substituents without dramatically altering binding affinity. However, while several substitutions of the pyrrolidin-2-one ring did not affect binding affinity, they had a dramatic effect on the residual activity of the enzyme-inhibitor-substrate complex (Tab. 1). For example, replacement of the pyrrolidin-2-one moiety of **1** (IC_50_ of 4 µM) with dimethylpropanamide in **1b** (IC_50_ of 3.6 µM) shifts the activity of the enzyme-inhibitor substrate complex from 1.4% to 61.7% (Tab. 1). Similar effects were observed for 4-methylaniline **1e** and 2-methoxyaniline derivative **1f** with IC_50_ of 8.5 and 10.7 µM respectively. Compound **1e** exhibits almost complete inhibition with only 3.1% DGC activity upon inhibitor saturation while **1f** functions as a partial inhibitor for DgcA with 59.4% activity remaining in the enzyme substrate inhibitor complex. These results suggest that **1f** binding converts the DgcA enzyme into a modified but still functional enzyme–substrate-inhibitor complex. Simultaneous binding of substrate and inhibitor indicates that **1** and its derivatives likely bind to a site distinct form the substrate binding site of DgcA.

**Table 1.**
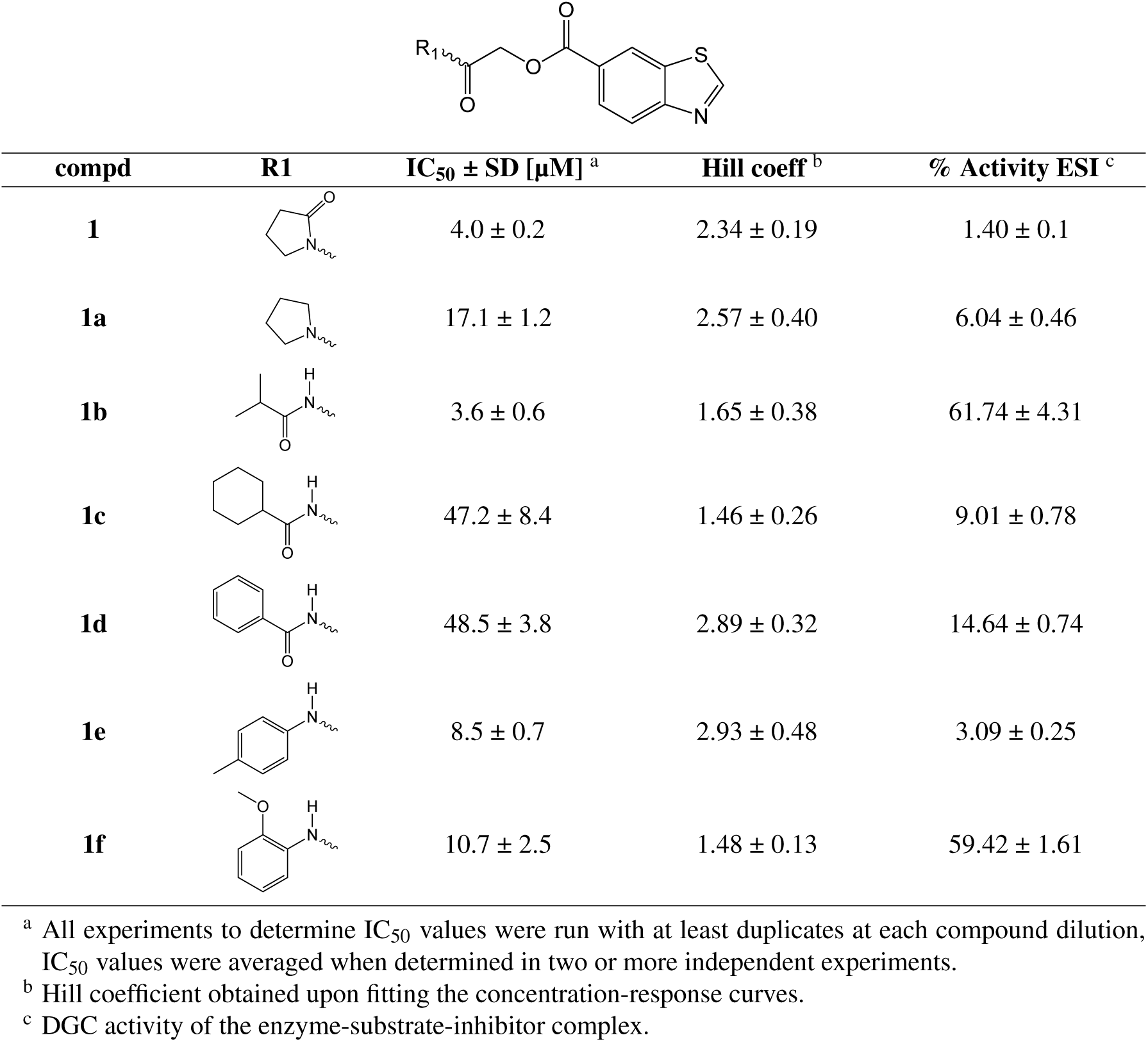
Exploration of pyrrolidin-2-one ring substitutions in the **1** scaffold.

| compd | R1 | IC<sub>50</sub> ± SD [μM]<sup>a</sup> | Hill coeff<sup>b</sup> | % Activity ESI<sup>c</sup> |
| --- | --- | --- | --- | --- |
| **1** |  | 4.0 ± 0.2 | 2.34 ± 0.19 | 1.40 ± 0.1 |
| **1a** |  | 17.1 ± 1.2 | 2.57 ± 0.40 | 6.04 ± 0.46 |
| **1b** |  | 3.6 ± 0.6 | 1.65 ± 0.38 | 61.74 ± 4.31 |
| **1c** |  | 47.2 ± 8.4 | 1.46 ± 0.26 | 9.01 ± 0.78 |
| **1d** |  | 48.5 ± 3.8 | 2.89 ± 0.32 | 14.64 ± 0.74 |
| **1e** |  | 8.5 ± 0.7 | 2.93 ± 0.48 | 3.09 ± 0.25 |
| **1f** |  | 10.7 ± 2.5 | 1.48 ± 0.13 | 59.42 ± 1.61 |

<sup>a</sup> All experiments to determine IC<sub>50</sub> values were run with at least duplicates at each compound dilution, IC<sub>50</sub> values were averaged when determined in two or more independent experiments.
<sup>b</sup> Hill coefficient obtained upon fitting the concentration-response curves.
<sup>c</sup> DGC activity of the enzyme-substrate-inhibitor complex.

### Effect of benzothiazole substitutions in scaffold 1 on binding affinity and modulation of DgcA

Next, we replaced the 1,3-benzothiazole-6-carboxylate moiety on **1** and chemically synthesized 12 derivatives containing alternative heterocycles and phenylsubstituents (materials and methods). We introduced various heterocycles with both 5- and 6-substituted carboxylates. In addition, we altered position and type of heteroatoms present in the heterocycles. For every derivative, we measured dose-response curves and maximal inhibition values using the FRET-based DGC assay. The inhibitory activity data and maximal inhibition values are summarized in Tab. 2. While compound **1i** in which the 1,3-benzothiazole-6-caboxylate was substituted with a 2-methyl-1,3-benzothiazole-5-carboxylate exhibited decreased inhibitory effects (IC_50_ 52.7 µM), we observed almost identical IC_50_ values for 1,3-benzoxazole-6-carboxylate **1g** and 2,3-dihydrobenzofuran 5-carboxylate derivative **1j** with 3.1 and 4.4 µM respectively (Tab. 2). However, **1g** and **1j** fail to effectively block DGC activity and DgcA remains partial active in presence of inhibitor (87.7% and 92.2%). These findings suggest that **1g** and **1j** both act as silent allosteric modulators (SAM) of DgcA (Tab. 2). Furthermore, we found, that substitution with indole-5-carboxylate **1k**, 1-methyl-indole-5-carboxylate **1l**, 1-methylbenzimidazole-5-carboxylate **1m**, isoquinoline-6-carboxylate **1n** and quinoxaline-6-carboxylate **1o** resulted in strongly reduced binding affinities towards DgcA but maintained their function as negative modulators. Similarly, alteration of the carboxylate side chain to 1,3-benzothiazole-5-carboxylate **1p** had a dramatic effect and decreased the binding affinity from 4 µM to 626 µM. These findings underline the isosteric importance of the 1,3-benzothiazole-6-carboxylate in mediating inhibitor potency of **1**.

**Table 2.**
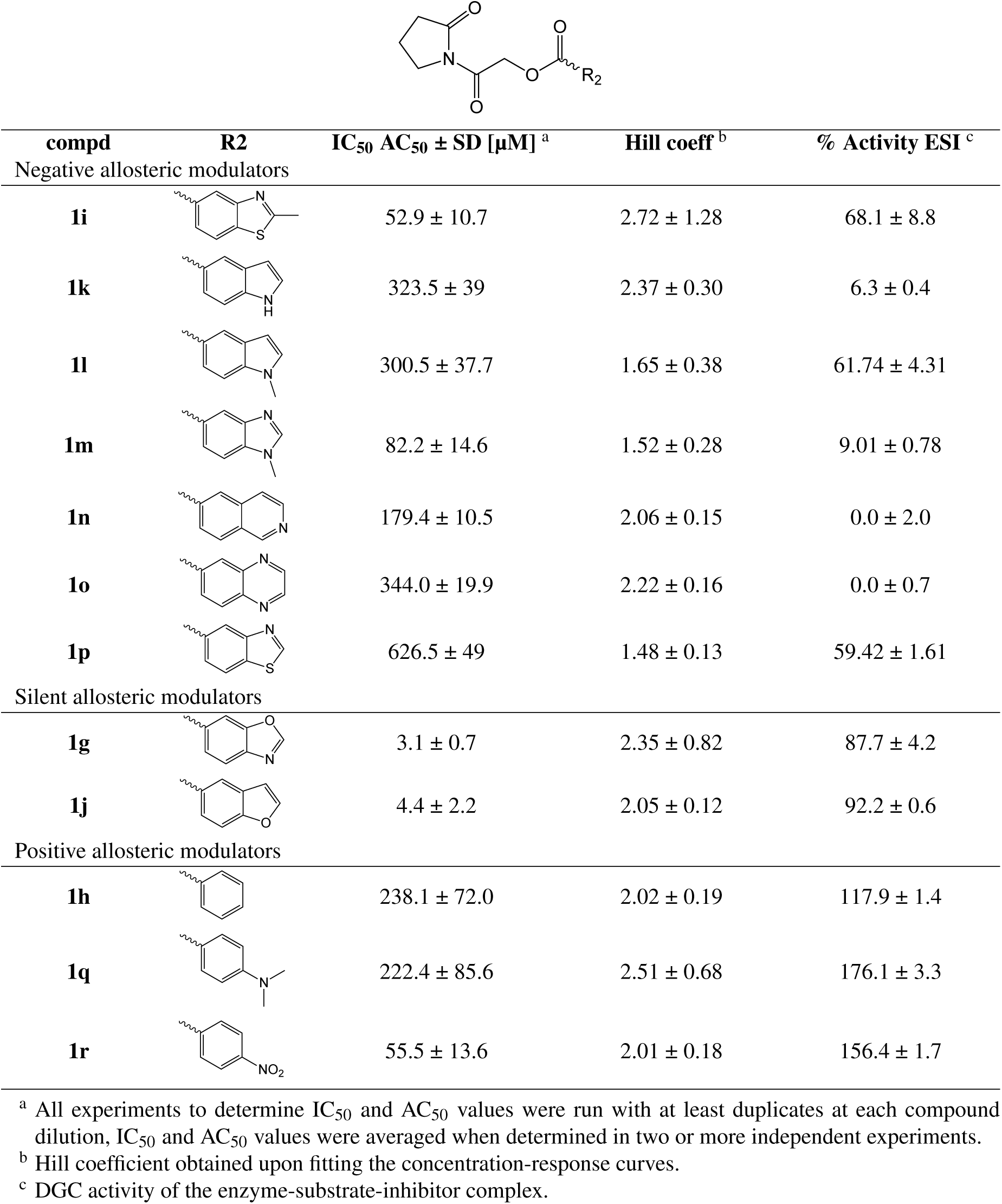
Substitution of 1,3-benzothiazole in scaffold **1** with alternative heterocycles and phenylsubstituents.

Next, we introduced benzoate **1h**, 4-(dimethylamino)benzoate **1q** and 4-nitrobenzoate **1r** (Tab. 2). Surprisingly, while introduction of these substituents resulted in rather moderate binding affinities, we observed that DgcA activity increases in presence of these compounds to activities higher than in absence of compounds (118%, 176% and 156%), suggesting that these compounds act as positive modulators of the DGC enzyme.

In the last group of derivatives, we investigated effects of linker modifications. The parental compound **1** contains an ester bond, which harbors the intrinsic risk of cleavage by cellular esterases. A nonhydrolizable amide analog would permit a much better stability to proteolysis by esterases. Therefore, we synthesized the nonhydrolizable amide derivative **1s** and tested its inhibitory properties against DgcA. We found that replacement of the ester bond with an amide increases IC_50_ from 4 µM to 31.6 µM and results in a partial inhibitor with 48% residual activity of the enzyme-inhibitor-substrate complex (Table S1). This suggests that differences in hydrogen bond forming properties or increased structural flexibility of the C-O-C bond might be important determinants for generating affinity and potency of the scaffold **1**. Similarly, introduction of a methyl group in the linker region **1t** or extension of the linker space between pyrrolidin-2-one and 1,3-benzothiazole-6-carboxylate **1u** completely abolished binding affinity, suggesting that the composition of the linker region as well as the relative orientation and distance between the pyrrolidin-2-one and benzothiazole moiety are critical determinants of binding affinity for DgcA.

While the parental compound **1** exhibits characteristics of a complete inhibitor, we created a panel of derivatives of **1** with diverse effects on DgcA activity. Several analogs of the scaffold **1** exhibit partial inhibition, some act as silent modulators or even function as positive modulators of DgcA. Thus, it seems likely that, rather than binding in a competitive mode, compounds of the scaffold **1** interact with an allosteric site of DgcA. The fact that structurally very similar compounds exert diverse effects on the activity of the enzyme-inhibitor-substrate complex suggests a model in which an allosteric binding site of DgcA is highly responsive to slight conformational changes. For example, substitution of a single sulfur atom in the benzothiazole moiety of **1** to oxygen in **1g** preserves binding affinity (IC_50_ of 4 and 3.1 µM respectively) but increases activity of the enzyme-inhibitor complex more than 60-fold from 1.4% to 87.7%. Similar effects are observed between **1** and **1b** where substitution of the pyrrolidin-2-one with dimethylpropanamide does not affect IC_50_ values but increases activity of the enzyme inhibitor complex from 1.4% to 62%. In conclusion, our structure activity studies on scaffold **1** suggest a model in which subtle substitutions within distal pyrrolidinone and benzothiazole moieties both affect affinity and determine if derivatives of the scaffold **1** act as negative, silent, or positive allosteric modulators of DgcA activity.

### Structure activity relationships of scaffold 2

Similar to several of the derivatives from **1**, the second scaffold 4-(2,5-dimethylphenoxy)-N-[4-(morpholin-4-yl)phenyl]butanamide **2** also acts as a partial inhibitor. Thus, we hypothesized that **2** might bind in a mechanism similar to the one observed for **1** and its derivatives **1a**–**u**. If both scaffolds bind to the same regulatory site of DgcA, it should be feasible to alter the modulatory effect of scaffold by exchanging the position and modification of the phenylsubstituents in **2** to generate derivatives with negative, silent, or even positive allosteric modulatory effects. We tested a set of 14 analogues of **2** by altering the phenylsubstituents on R1 and R2 and keeping the butanamide linker constant (Tab.1). The parental scaffold **2** harbors two methyl substituents on R1 and one 4-morpholino substitution at R2 (Tab. 3). We found that removal of all phenylsubstituents in **2c** does not significantly alter binding affinity (10.6 µM versus 6.4 µM) or activity of the enzyme modulator bound complex (38.6% versus 31.4%). For compound **2d**, harboring a single 4-methyl substitution, we observed a binding affinity of 207 µM. For the 2-methyl substituted derivative **2e**, we determined an IC_50_ of 2.2 µM. The steric load of a single methyl group at the para position barely accounts for a 100-fold difference in binding affinity between **2d** and **2e**. Similar results were observed for alterations in the methyl substitution pattern in compound **2a**. Movement of a methylgroup from 5- to 4-position at R1 preserved affinity at 11.7 µM meanwhile increased activity of the enzyme inhibitor complex from 31.4% to 69.8%. However, removing either one of the methyl substitutions in **2a** resulted in derivatives with differences in binding affinity.

**Table 3.**
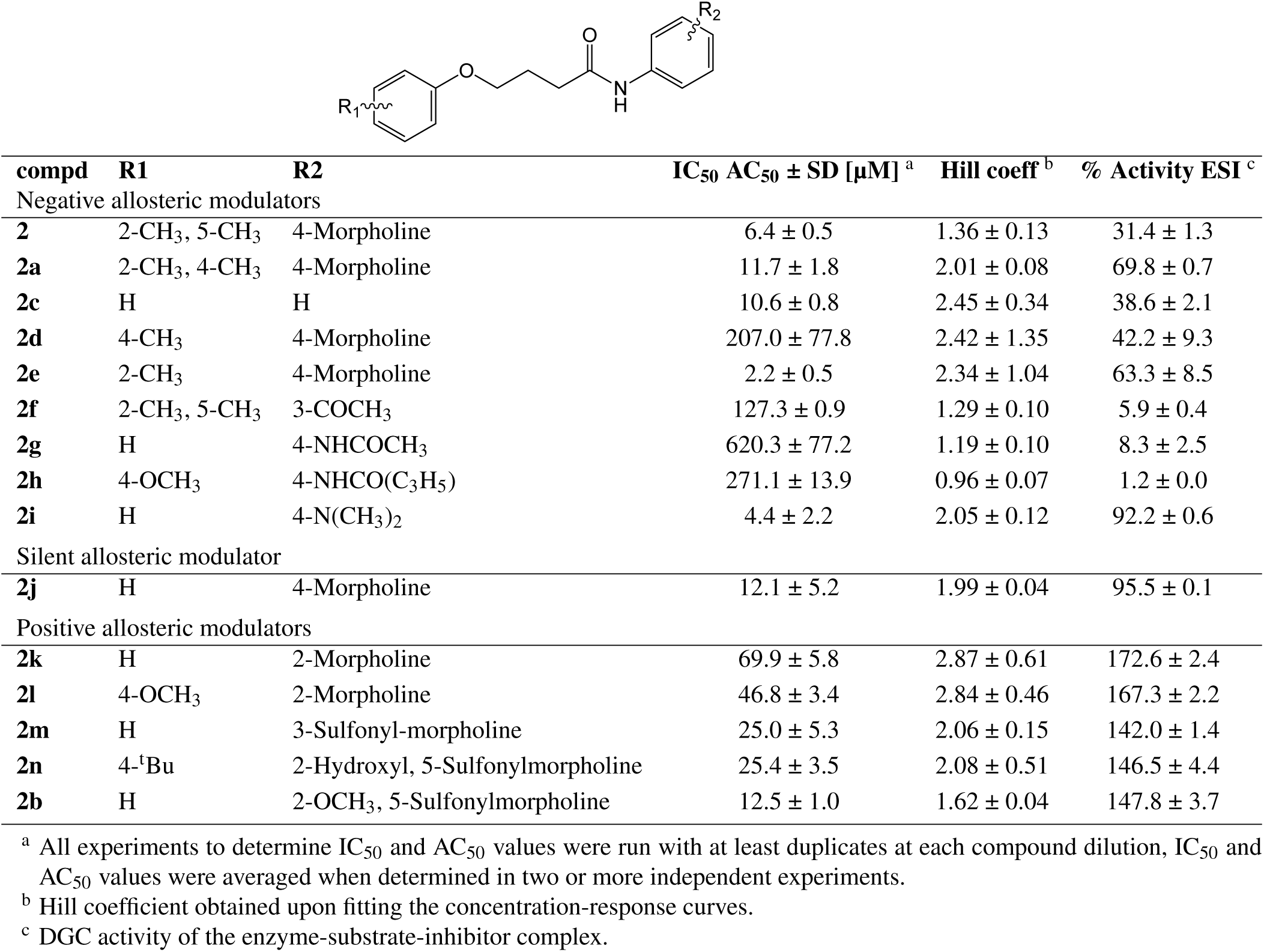
Exploration of phenylsubstituents in **2** and their modulatory effect on DGC activity.

We further expanded the spectrum of negative allosteric modulators by introducing at R2 substitutions with 3-acetyl **2f**, 4-acetamide **2g**, 4-cyclopropanecarboxamide **2h** and 4-dimethylamino **2i**. With exception of the dimethylamino derivative **2i**, which retained 71.1% DGC activity, all compounds have inhibitor bound activities smaller than 8.5%. Substitutions patterns differ in the group of negative allosteric modulators of the scaffold **2** with respect to steric demand and hydrogen-bonding properties, however, most of them contain para-substitutions at R2. We investigated the modulatory effects of ortho and metha-substitutions at R2 and introduced either at the ortho- position morpholine **2k**,**l** or at the meta-position sulfonylmorpholine substitutions **2b**,**m**,**n**. Upon measuring dose-response curves and maximal inhibition values using the FRET based DGC assay, we observed for all meta- and ortho-substituted derivatives **2b**,**k**–**n** increasing DGC activity between 142% and 172%, suggesting that these compounds act as positive modulators of the DGC enzyme (Fig. 3D and Tab. 3).

**Fig 3.**
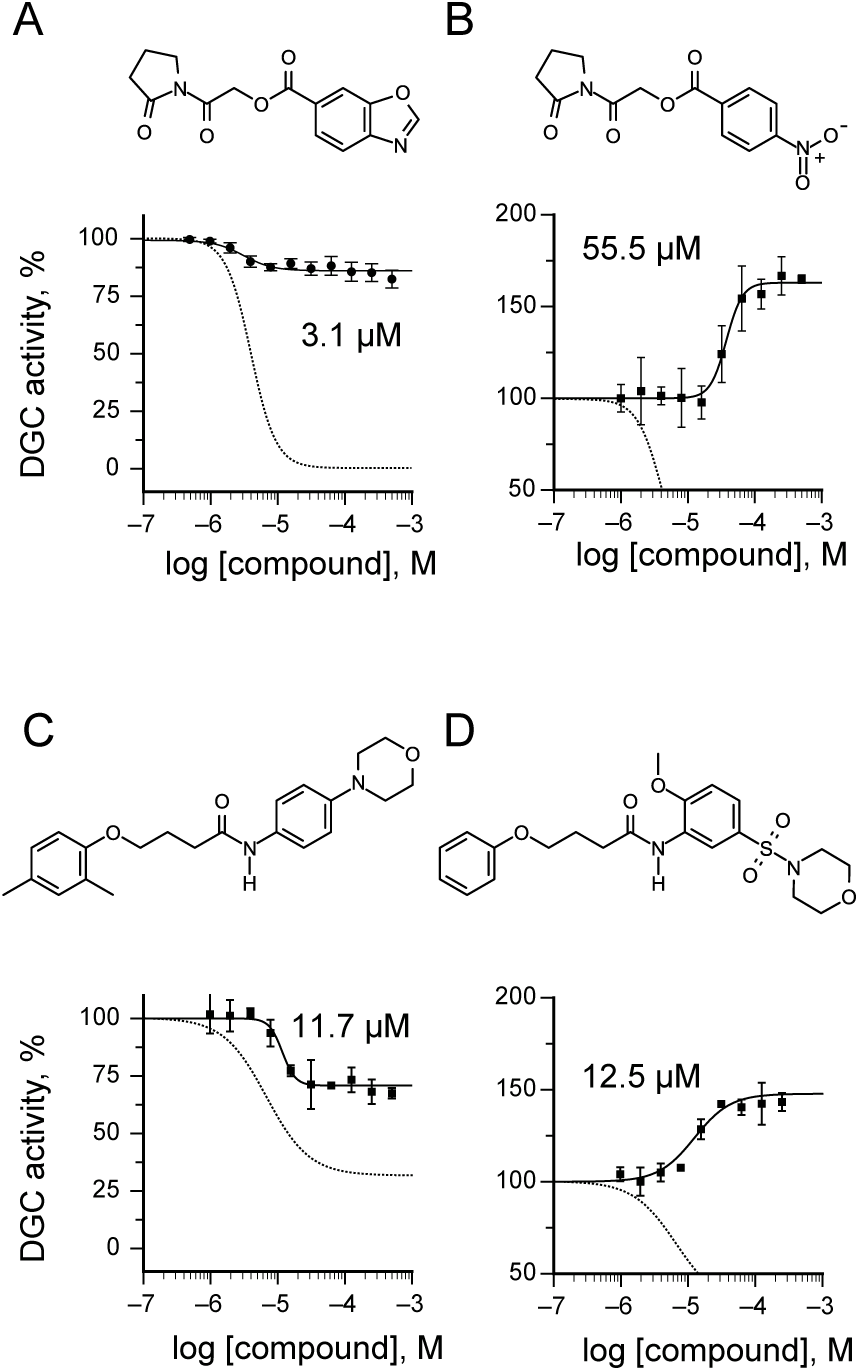
Examples of silent (SAM), negative (NAM) and positive (PAM) allosteric modulators of the DGC enzyme DgcA. (**A**) Structure and concentration response curve of the 1,3-benzoxazole-6-carboxylate derivative **1g**, a silent allosteric regulator of DgcA. (**B**) Representative example of an allosteric activator of the scaffold **1**. The 4-nitrobenzoic acid derivative of **1h** acts as a positive allosteric regulator with AC50 of 55.5 µM. The concentration response curves of the parental 1,3-benzothiazole-6-carboxylate **1** are shown as dashed lines in panels A and B. (**C**) Structure and concentration response curve of **2a**. Alteration of the methyl substituents from 2,5 to 2,4 in the **2** scaffold increases residual activity of the enzyme-substrate-inhibitor complex from 31.4% in **2** (dashed line) to 69.8% in **2a** (closed circles). (**D**) The 2-methoxy, 5-sulfonylmorpholine derivative **2b** is a positive allosteric regulator with AC50 of 12.5 µM. Concentration response curves of the parental **2** are shown as dashed lines in panels C and D.

### Mechanism of inhibition for 1 and 2

Our results suggest that both scaffolds **1** and **2** are able to bind to the DgcA enzyme at a site distinct from the GTP binding pocket. Although, it may be possible for such mixed-type modulators to bind at the active site, allosteric modulation generally results from compounds binding to different sites. An attractive candidate for such an allosteric binding site is the product inhibition site (I-site) of DgcA. Product inhibition by c-di-GMP is a general regulatory mechanism of many DGC enzymes and requires a RXXD motif which forms a turn at the end of a short five-amino acid linker that directly connects the I-site with the conserved catalytic A-site residues, GG(D/E)EF(9).

Previous molecular simulations on the DGC PleD in presence and absence of c-di-GMP revealed a significant decrease in mobility for both I- and A-site residues upon complex formation (9). This suggests that dynamic coupling between the two sites via the short connecting beta-strand and a balance-like movement could be responsible for direct information transfer. To investigate whether allosteric binding of scaffolds **1** and **2** affect substrate binding or maximal reaction rate of DgcA at the active site, we performed DGC inhibition assays in presence of increasing concentrations of substrate GTP (Fig. 4 and Tables S2, S3). We found that binding of **1** affects only the Km for GTP but does not impair maximal velocity of enzyme catalysis (Fig 4A,B and Table S2). This suggests that binding of **1** at the allosteric site induces conformational changes at the active site resulting in reduced affinity for substrate binding but maintains the three-dimensional conformation of catalytic residues. In contrast, we observed for **2** a decrease in affinity for substrate binding as well as a reduction in maximal rate of the DGC reaction (Fig 4A,B and Table S3). Thus, binding of **2** likely destabilizes the divalent Mg2+ carboxyl complex that coordinates and activates the triphosphate moiety of GTP as well as affects the conformation of additional residues at the GTP binding site.

**Fig 4.**
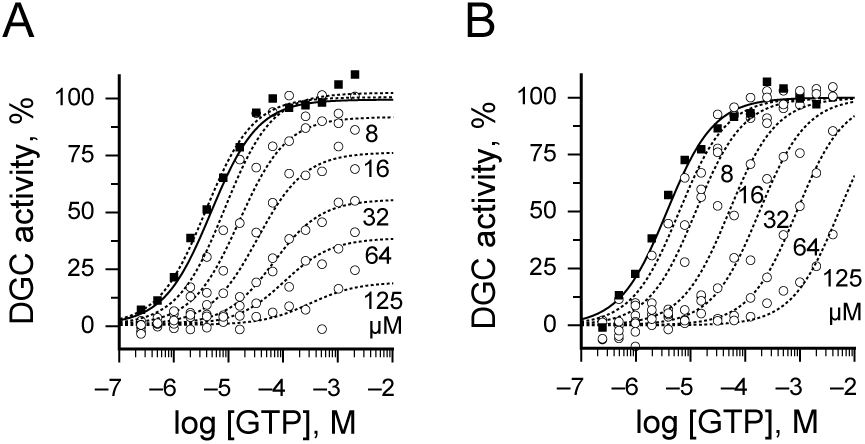
Effect of **1** and **2** on substrate binding and maximal reaction rate of DgcA. Corresponding plot of the initial velocity versus GTP concentration. (**A**) Presence of **1** decreases affinity for substrate binding and maximal rate of DGC reaction (Table S3). (**B**) In contrast to **1**, binding of **2** affects only the Km for GTP but does not affect maximal velocity (Table S3).

## Discussion

Due to their importance in governing the amount and spatiotemporal distribution of c-di-GMP, DGC enzymes are considered important targets for control of biofilm formation and antibiotic persistence phenotypes. Our results demonstrate that fluorescence protein based FRET biosensors can be employed as powerful analytical tools to detect c-di-GMP in the nanomolar range and monitor performance of diguanylate cyclase enzyme reactions in a 384 well format in the low nanomolar range. With increasing numbers of assays available that monitor fluctuations in cellular c-di-GMP levels (10, 15–17), we can begin to identify effective approaches to discover potent small molecule modulators of c-di-GMP signaling and monitor their phenotypic effects and influence on cellular c-di-GMP signaling patterns. In addition to c-di-GMP, many other cyclic nucleotide second messengers such as cAMP, cGMP, c-di-AMP and cGAMP currently lack HTS capable analytical methods that allow in situ detection of their levels during enzymatic assays or within complex biological samples. In the future, similar FRET-based biosensors could provide an interesting alternative to current existing assays. Often, receptor proteins are known to recognize their cognate signaling molecule with high specificity. Insertions of these receptor domains between a FRET pair of fluorescent proteins will lead to novel biosensor designs that detect their cognate signaling molecule with high specificity. Several strategies for the chemical synthesis of c-di-GMP derivatives as potential inhibitors targeting DGC have been reported (18, 19) and a number of small molecule inhibitors of DGC activity has been previously described (20–24). While these compounds were valuable research tools, their potency and pharmaceutical properties were insufficient to determine the impact of DGC inhibition in vivo. Originally our HT screen was designed to discover inhibitors of diguanylate cyclases. However, our subsequent structure activity studies unraveled a panel of 44 compounds that exert surprising variations in modulatory effects on DgcA activity. Among them is one complete negative allosteric modulator **1** with IC_50_ of 4 µM, two partial negative regulators **1b**,**c** with 3.6 and 2.2 µM, two silent allosteric modulators **1g**,**j** with IC_50_ of 3.1 and 4.4 µM and one positive allosteric modulator of DgcA activity **2b** with AC_50_ of 12.5 µM.

In terms of activities, we discovered a remarkable diversity between compounds with even small chemical substitutions on side chains reverting effects between inhibition and DGC activation. These results emphasize the importance of dynamic conformational changes rather than the static three-dimensional structure for the biological function of DGC modulators. For example within scaffold **1**, substitution of a single sulfur atom in benzothiazole with oxygen increases activity of the modulator bound complex over 60-fold while maintaining binding affinity at 4 µM. Within the **2** scaffold, shifting a single para-substitution to ortho-position, converted the silent modulator **2j** into an activator **2k**.While previously the I-site has been reported as an inhibitory site (8, 10) our analysis reveals that DGC enzymes are subjected to additional layers of allosteric regulation. Rather than inhibiting, we demonstrate that several synthetic small molecules are able to allosterically activate DgcA. It is interesting to speculate that other endogenous molecules or protein domains might adopt a similar mechanism of action and function as endogenous allosteric activators of DGC enzymes coupling DGC activity to distinct cellular signaling inputs. In fact several GGDEF domain containing proteins, despite harboring all critical conserved active site residues, fail to exhibit DGC activity in biochemical assays (25, 26). Such enzymes might exhibit conditional DGC activity in the presence of specific endogenous activators.

In terms of drug discovery, our results demonstrate the complex interplay between synthetic small molecules and the regulatory mechanism that drives DGC activation. In many prokaryotic model systems, c-di-GMP mediated biofilm formation is regulated by multiple DGC enzymes in parallel. Thus our studies underline the importance for in detail mechanism of action studies of candidate DGC inhibitors prior to their phenotypic evaluation. In the future, it will be interesting to investigate whether distinct fragments of the compounds identified will act as general DGC modulators or exhibit specificity towards distinct DGC subtypes. Compounds that act specifically on distinct enzymes may serve as exciting chemical genetics tools for the dissection of DGC-networks while inhibitors with generalized effects of DGC activity will have the potential to serve as lead compounds for the development of biofilm modulators and other c-di-GMP mediated phenotypes involved in bacterial pathogenesis.

## METHODS

Supplementary Methods includes descriptions of (i) compounds, storage of proteins and reagents (ii) methods for protein expression, storage of protein and reagents, diguanylate cyclase enzyme assay and, (iii) HT screening, and data analysis and chemical synthesis procedures.

## ACKNOWLEDGEMENTS

We thank Jonathan Venetz for comments on the manuscript. Compounds for the high-throughput screening were provided by the national screening laboratory for the regional centers of excellence in biodefense and emerging infectious diseases (NIAID U54 AI057159). We thank Su Chaing and the staff at the NERCE national screening laboratory (NSRB) for assistance in performing the high-throughput screening and post screen analysis.

## AUTHOR CONTRIBUTIONS

M.C. and D.H.K designed the experiments, developed FRET assay and performed HT screening, K.C.O and T.K. performed chemical synthesis and validated hits for synthetic tractability, M.C, C.K. and B.K performed does response measurements, MC. D.H.K, B.C and S.I.M. analyzed data, M.C, B.C, K.C.O, D.H.K and S.I.M. wrote the manuscript.

## Supplementary 1

### Methods

#### Compound library

Compounds for screening were provided by the national screening laboratory NSRB, in collaboration with the ICCB-Longwood screening facility located at Harvard Medical School. The chemical library screened is a subset of a compilation of commercially available small molecule chemical libraries maintained at the ICCB. The compounds have been prescreened for drug-like properties (Lipinski’s rules)(27). Stock libraries were stored at a concentration of 5 mg/mL in DMSO at –80 °C in 384-well format, with at least two empty columns on each plate to allow for on-plate controls. Libraries (number of molecules) from the following companies were evaluated: Asinex (12377; Winston-Salem, NC), Enamine (11264; Kiev, Ukraine) and Life Chemicals (3893; Burlington, Canada).

#### Compounds used for dose-response (IC_50_) assays

For confirmatory assays and dose-response (IC_50_) assays the following compounds were purchased from Enamine, Ltd. (www.enamine.net, Kyiv, Ukraine): **1** (T5267517), **1a** (T5382986), **1b** (T5423966), **1c** (T5354248), **1d** (T5283214), **1e** (T5290508), **1f** (T5841885), **2g** (T5257569), **2h** (T6309238), **2i** (T5401340), **2k** (T5558380), **2l** (T5555070), **2m** (T5577173), **2n** (T5553390), **2b** (T5577172), **2o** (T6283377), **2p** (T0520-5955), **3** (T5580955), **4** (T5411955), from Asinex Corporation (www.asinex.com, Winston-Salem, NC 27101, USA); **MDC2** (BAS 05165102), **5** (BAS 01811924), **MDC7** (BAS 14051565), **2c** (BAS 01833594), **2d** (BAS 03310955), **2e** (BAS 03079336), **2j** (BAS 01835812), from ABI Chem (www.abichem.com, Munich, Germany); **6** (AC1N893M), **2a** (AC1LV267) and from Molport Corporation (www.molport.com, Riga, Latvia); **2f** (MolPort-002-121-569).

#### Screening equipment

Multi-well liquid dispensing for setting up assays was performed using the liquid handler PlateMate Plus (Hamilton) and Microflo plate dispenser (Biotek). High-throughput Screening, DGC enzyme inhibition reactions, and dose-response (IC_50_) assays were performed in Corning Low Volume 384 Well Black Flat Bottom Polystyrene Microplates (Product #3821BC) in a final volume of 20 µl per well. Change in fluorescence ratio (535/470nm) was measured in Envision 2104 fluorescence plate reader (Perkin Elmer) to monitor for c-di-GMP production. The plate reader was equipped with CFP excitation filter (Ex 430nm, BW 24nm and Tmin 90%) a dichroic mirror module (D450/D505 nm), emission filters for YFP (CWL 535nm, BW 30nm and Tmin 90%) and CFP (CWL 470nm, BW 24nm and Tmin=90%) and dual PMT detectors for recording CFP and YFP signals simultaneously. Measurement mode was bi-directional by rows (excitation and emission light set to 100% and number of flashes to 10) using dual detector mode with a PMT gain of 383 (535 nm) and 572 (470 nm) and a measurement height of 7 mm above the plate height.

#### Protein expression

*E. coli* BL21 (DE3) strain BC1058 (9) containing the constitutive active *Caulobacter crescentus* DGC (CC3285) cloned into pET42b plasmid was used for recombinant expression of the DGC enzyme. DgcA protein was overexpressed and His-tag purified according to (9). E. coli BL21 (DE3) strain containing pET15b::mCPet-12AA-mYPet plasmid was used for recombinant expression of the c-di-GMP biosensor. The expression vector used for protein purification of c-di-GMP biosensors was a modified variant of the pET15b::mCPet-12AA-mYPet (28) containing an N-terminal polyhistidine tag for affinity purification, monomerizing mutations in CYPet (A206K) and YPet (A206K) and a multiple cloning site in its 12 amino acid linker region between the mYPet and mCYPet FRET pair. Two restriction sites (SpeI and KpnI) were introduced by PCR into the 5’ and 3’ ends of a codon optimized version of Salmonella enterica serovar Typhimurium ycgR (STM1798) gene and the products were cloned in-frame into the SpeI and KpnI sites of pET15b::mCPet-12AA-mYPet (10). YcgR biosenso was overexpressed and His-tag purified according to (9).

#### Storage of Protein and Reagents

All proteins used in our assays were purified using affinity chromatography followed by gel filtration. These enzymes were kept at a low concentration (less than 0.1mg/mL to prevent aggregation) in a solution containing 20% glycerol, protease inhibitors and b-mercapto ethanol. The DGC enzyme is active for at least 2 weeks at 4°C and >90% active for at least 48 hours at room temperature. The YcgR biosensor can be stably kept for weeks in the dark at 4°C. For long-term storage the biosensor was stored at −80 °C, in small aliquots containing 20% glycerol and tris buffer (10mM) pH 8 and 250mM NaCl. We have tested the biosensor that has been frozen for more than 1 year for stability and it did not show any significant loss of activity. C-di-GMP is kept at −20°C for long-term storage as a liophylized powder or solution. Our current stocks are made in-house biosynthetically using a constitutive active derivative of DgcA enzyme (9) and purified using HPLC. GTP stock solutions are prepared just before the assay or screening as it will hydrolyze over time.

#### Diguanylate cyclase assay

DGC reactions were performed at 25°C with 20 nM purified hexahistidine-tagged DgcA and 50 nM hexahistidine-tagged mCPet-YcgR-mYPet biosensor to monitor change in fluorescence upon c-di-GMP production. The DGC reaction buffer contained 250 mM NaCl, 25 mM Tris-Cl, pH 8.0 and 20 mM MgCl2. For screening assays, the reaction mixture was preincubated with 200 nL compound (5 mg/mL) in DMSO and fluorescence emission was measured prior addition of YcgR biosensor, then again after addition of GTP (t = 0 min) and 3 h post GTP addition as the end-point measurement. For IC_50_ inhibition assays, the protein was pre-incubated with different concentrations of compounds dissolved in DMSO (1–100 µM, 1% final DMSO concentration) for 2 min at 25 °C before GTP substrate was added to a final concentration of 20 µM. During Screening, fluorescence emission ratio was measured prior addition of the YcgR biosensor to monitor background fluo-rescence at 470 and 535 nm. Plates were directly measured after addition of 20 µM GTP (t =0 min) and then after 3h incubation at RT. During secondary assays and IC_50_ inhibition assays change in fluorescence ration was measured at time intervals of 1 min. We determined the plate uniformity, signal variability and repeatability assessment using NCGC guidelines. The data was analyzed using the Excel file supplied by the NCGC assay validation website. We conducted the “plate uniformity assay” recommended by NCGC for CV and Z-factor calculation for measuring plate-to-plate variation as well as day-to-day variation. The average Z-factor for 3 plates, read on 2 consecutive days was 0.71 and the CV for high, medium and low signal were well below 5%.

#### Data Analysis

Data analysis was performed in Excel 97. All primary screening was performed at 50 µg/mL compound concentrations. The fluorescence emission ration 535/470 nM of the YcgR biosensor in absence of c-di-GMP was used as the positive control for DgcA inhibition and biosensor in the presence of 5 µM c-di-GMP served as the negative control. For a compound to be considered a hit, the following criteria had to be met: (1) compounds with c-di-GMP levels below 300 nM after 3h incubation with DgcA and 20 µM GTP were considered strong hits; (2) duplicates must be consistent with difference in final c-di-GMP levels <100 nM; and (3) background fluorescence prior addition of the FRET biosensor must not exceed 7% of the FRET signal in CFP and YFP emission channels. Total fluorescence was used as a quality control criterion to identify inherently fluorescent molecules. From the 27502 compounds screened in duplicates, 1029 (3.7%) were removed from the analysis due to auto fluorescence. 49 compounds (0.19%) exhibited c-di-GMP levels below 300 nM after 3h incubation with DgcA enzyme. Out of these, 18 compounds were tested in secondary assays. 7 had potencies below 50 µM and 3 compounds below 10 µM.

The FRET fluorescence emission ratio was defined as:

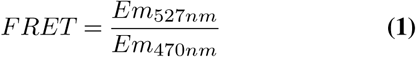

Θ, The fraction of YcgR biosensor in the c-di-GMP bound state was calculated according to:

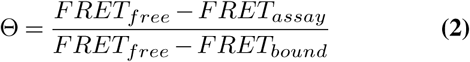

Where *FRET*_*free*_ fluorescence emission ratio 535/470nm of the free biosensor, *FRET*_*bound*_ fluorescence emission ratio 535/470 nm of YcgR with c-di-GMP bound and *FRET*_*assay*_ fluorescence emission ratio 535/470 nm of the YcgR biosensor during the assay.

The c-di-GMP concentration (in nM) was calculated according to:

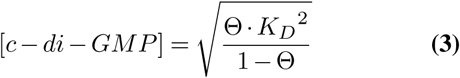

With *K*_*D*_ = 198*nM*, c-di-GMP dissociation constant of the YcgR biosensor at 25°C Initial velocity *V*_*o*_ was determined by plotting c-di-GMP concentration versus time and by fitting the curve according to Michaelis-Menten kinetics. The doseresponse curves of Percent Activity were fit in ProFit 5.6.7.

IC_50_ inhibition constants were fitted with the following function:

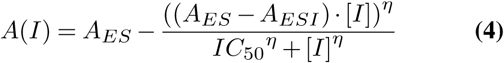

Where A(I) = relative DGC enzymatic activity as a function of inhibitor concentration, *A*_*ES*_ = relative activity in absence of inhibitor, *A*_*ESI*_ = relative residual activity under inhibitor saturating conditions. *IC*_50_ = half maximal inhibitory concentration of inhibitor, [*I*] concentration of inhibitor and η = hill coefficient.

AC50 constants were fitted with the following function:

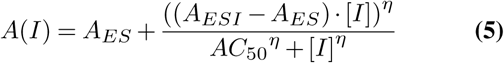

Where A(I) = relative DGC enzymatic activity as a function of activator concentration, *A*_*ES*_ = relative activity in absence of activator, *A*_*ESI*_ = relative activity under activator saturating conditions. *AC*_50_ = half maximal activating concentration of activator, [*I*] concentration of activator and *η*= hill coefficient.

All hit validation and high-throughput screening procedures were carried out according to the general guidelines as published on the U.S. National Chemical Genomics Center web site (NCGC Assay Guidance Manual and High-throughput Assay Guidance criteria (http://www.ncbi.nlm.nih.gov/books/NBK53196/).

## Chemistry

### General Methods

All reactions were run under an atmosphere of dry nitrogen. Reagents and solvents were obtained in the highest available purity and used without further purification unless indicated. ^1^H NMR and ^13^C NMR spectra were obtained on a 500 MHz (Bruker AV500) instrument. Identity of the compounds was confirmed by mass spectrometry. The compound solution was infused into the electrospray ionization source operating in positive or negative ion mode. Low resolution spectra were obtained on the Esquire LC ion trap mass spectrometer (Bruker Daltonics, Billerica, MA). Accurate mass measurements were performed on the APEX Qe 47 Fourier transform ion cyclotron resonance mass spectrometer (Bruker Daltonics, Billerica, MA). Normal phase silica gel purifications were done using a Biotage SP4 instrument using the cartridges supplied by Biotage. RP-HPLC was done on a Varian instrument equipped with a diode array ultraviolet detector. For preparative reverse phase chromatography a 10 x 250 mm C18 5µ column at a flow rate of 4.6 mL/min was used; for analytical reverse phase chromatography a 4.6 x 250 mm C18 5µ column at a flow rate of 1 mL/min was used. Ultraviolet detection was at 215 and either 254 or 360 nm. Unless otherwise specified, buffer A was 0.05% TFA in H_2_O, buffer B was 0.05% TFA in acetonitrile. Thin layer chromatography was done using 0.2 mm polygram SIL G/UV plates (Alltech, Deerfield, IL), developed using mobile phases of varying compositions of ethyl acetate/hexane, MeOH/CH_2_Cl_2_, or MeOH/CHCl_3_, and visualized by UV light supplemented by vanillin, ninhydrin, and other solution stains where appropriate.

**Fig. S1.**
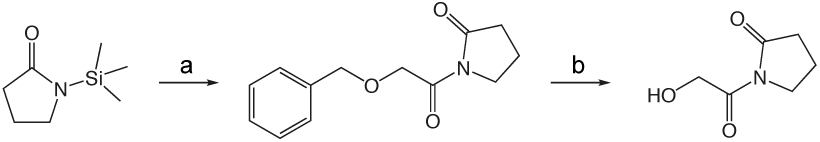

**Scheme 1. Synthesis of 1v.** a). benzyloxyacetyl chloride, ether 0°C - r.t. 1.5h. b). Pd-C, EtOH, DMF, H_2_

**Fig. S1a.**
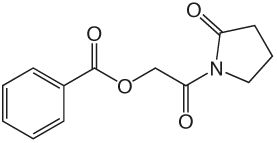

**1w** 1-(2-(benzyloxy)acetyl)pyrrolidin-2-one

### 1w

To a solution of benzyloxyacetylchloride (5.3 mL, 33.59 mmol) in 55 mL ether at 0 °C was added 1-TMS-2-pyrrolidinone (5.9 mL, 36.95 mmol) in 55 mL ether dropwise. The ice bath was removed and the reaction mixture stirred at room temperature 1.5 h, then concentrated in vacuo. The crude product was purified via silica gel chromatography using a gradient from 30 to 60% ethyl acetate in hexanes tom give **1w** (7.425 g, 31.87 mmol). ^1^H NMR (500 MHz, CDCl_3_, δ): 1.92-2.08 (m, 2H), 2.48 (td, J = 8.0 Hz, 3.6 Hz, 2H), 3.75 (td, J = 7.0 Hz, 3.7 Hz, 2H), 4.61 (s, 4H), 7.15-7.28 (m, 1H), 7.29 (t, J = 6.6 Hz, 2H), 7.32-7.49 (m, 2H). ^13^C NMR (500 MHz, CDCl_3_, δ): 17.74, 33.00, 44.78, 70.83, 73.21, 127.79, 127.94, 128.37, 137.53, 171.16, 175.70. MS m/z 256.2 [M + Na]^+^.

**Fig. S1b.**
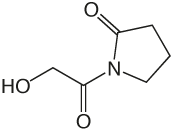

**1v** 1-(2-(benzyloxy)acetyl)pyrrolidin-2-one

### 1v

In a flame dried flask, 200 mL EtOH was bubbled with argon 10 min. 10% palladium on carbon (7.425 g) was added and the suspension bubbled with argon 10 min, then **1w** (7.425 g, 31.87 mmol) in 100 mL and 33 mL DMF added, bubbled with argon 10 min then bubbled with H_2_ until completion. The solution was bubbled with argon 10 min, filtered through celite and concentrated in vacuo to give **1v** (4.436 g, 31.02 mmol). ^1^H NMR (500 MHz, CDCl_3_, δ): 2.05-2.20 (m, 2H), 2.58 (t, J = 8.1 Hz, 2H), 3.83 (t, J = 7.2 Hz, 2H), 4.62 (s, 2H). ^13^C NMR (500 MHz, CDCl_3_, δ): 17.91, 32.88, 44.93, 64.17, 174.30, 175.92. MS m/z 166.1 [M + Na]^+^. HRMS (m/z): [M + Na + CH_3_OH]+1 calcd for C_7_H_13_NNaO_4_, 198.0737; found 198.0736.

### 1i (General Method A.)

To a solution of **1v** (49 mg, 0.34 mmol) in 2 mL CH_2_Cl_2_ and 0.5 mL DMF was added

### Scheme 2. General method A

a). 5-benzothiazole-carboxylic acid, EDCI, DMAP, CH_2_Cl_2_, DMF r.t. overnight

**Fig. S2.**
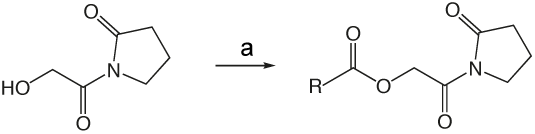

### 1i

**Fig. S2a.**
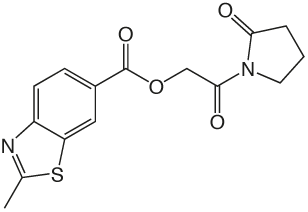

2-oxo-2-(2-oxopyrrolidin-1-yl)ethyl 2-methylbenzo[d]thiazole-6-carboxylate

EDCI (131 mg, 0.44 mmol), DMAP (3 mg, 0.027 mmol), and 2-methyl-1,3-benzothiazole-6-carboxylic acid (55 mg, 0.31mmol). The reaction mixture stirred at room temperature overnight then was partitioned between CH_2_Cl_2_ and saturated NaHCO_3_ solution. The organic layer was washed with brine, dried over Na2SO4, filtered, and concentrated in vacuo. The crude product was purified via silica gel chromatography using a gradient from 0 to 3% MeOH in CH_2_Cl_2_ to give **1i** (62 mg, 0.19 mmol). ^1^H NMR (500 MHz, CDCl_3_, δ): 2.08-2.21 (m, 2H), 2.62 (t, J = 8.1 Hz, 2H), 2.86 (s, 3H), 3.83 (t, J = 7.2 Hz, 2H), 5.42 (s, 2H), 7.98 (d, J = 8.6 Hz, 1H), 8.18 (dd, J = 8.6 Hz, 1.7 Hz, 1H), 8.62 (d, J = 1.3 Hz, 1H). ^13^C NMR (500 MHz, CDCl_3_, δ): 17.89, 20.46, 32.91, 44.87, 65.01, 122.16, 124.06, 125.84, 127.55, 135.65, 156.58, 165.70, 167.92, 171.06, 176.09. MS m/z 341.5 [M + Na]^+^. HRMS (m/z): [M + Na]^+^ calcd for C_15_H_14_N_2_NaO_4_S, 341.0566; found 341.0569.

**Fig. S2b.**
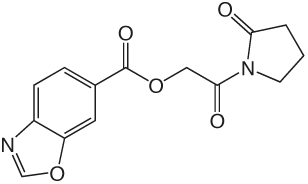

**1g**2-oxo-2-(2-oxopyrrolidin-1-yl)ethyl benzo[d]oxazole-6-carboxylate

### 1g

was prepared according to the procedure in General Method A from **1v** and benzo(d)oxazole-6-carboxylic acid on a 0.30 mmol scale. The crude product was purified five times via silica gel chromatography using a gradient from 0 to 5% MeOH in CH_2_Cl_2_ to give **1g** (25 mg, 0.087 mmol). ^1^H NMR (500 MHz, CDCl_3_, δ): 2.08-2.26 (m, 2H), 2.66 (t, J = 8.1 Hz, 2H), 3.87 (t, J = 7.2 Hz, 2H), 5.45 (s, 2H), 7.86 (d, J = 8.3 Hz, 1H), 8.19 (d, J = 8.3 Hz, 1H), 8.26 (s, 1H), 8.38 (s, 1H). ^13^C NMR (500 MHz, CDCl_3_, δ): 17.92, 32.93, 44.89, 65.13, 113.21, 120.40, 126.67, 127.15, 144.15, 149.65, 155.06, 165.56, 167.86, 176.13. MS m/z 311.2 [M + Na]^+^. HRMS (m/z): [M + Na]^+^ calcd for C_14_H_12_N_2_NaO_5_, 311.0638; found 311.0644.

1j 2-Oxo-2-(2-oxopyrrolidin-1-yl)ethyl benzofuran-5-carboxylate

### 1j

**Fig. S2c.**
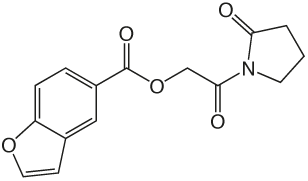

was prepared according to the procedure in General Method A from **1v** and 1-benzofuran-5-carboxylic acid on a 0.38 mmol scale. The crude product was purified via silica gel chromatography using a gradient from 0 to 5% MeOH in CH_2_Cl_2_ to give **1j** (23.6 mg, 0.082 mmol). ^1^H NMR (500 MHz, CDCl_3_, δ): 2.09-2.27 (m, 2H), 2.66 (t, J = 8.1 Hz, 2H), 3.87 (t, J = 7.2 Hz, 2H), 5.45 (s, 2H), 6.87 (d, J = 1.2 Hz, 1H), 7.57 (d, J = 8.7 Hz, 1H), 7.72 (d, J = 2.0 Hz, 1H), 8.12 (dd, J = 8.7 Hz, 1.4 Hz, 1H), 8.45 (s, 1H). ^13^C NMR (500 MHz, CDCl_3_, δ): 17.92, 32.97, 44.90, 64.87, 107.17, 111.37, 124.25, 124.43, 126.44, 127.49, 146.31, 157.71, 166.30, 168.17, 176.09. MS m/z 310.3 [M + Na]^+^. HRMS (m/z): [M + Na]^+^ calcd for C_15_H_13_NNaO_5_, 310.0686; found 310.0690.

**Fig. S2d.**
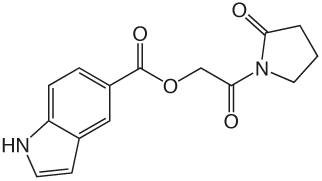

**1k** 2-oxo-2-(2-oxopyrrolidin-1-yl)ethyl 1H-indole-5-carboxylate

### 1k

was was prepared according to the procedure in General Method A from **1v** and indole-5-carboxylic acid on a 0.29 mmol scale. The crude product was purified four times via silica gel chromatography using a gradient from 10 to 50% ethyl acetate in hexanes to give **1k** (8.2 mg, 0.029 mmol). ^1^H NMR (500 MHz, DMF-d7, δ): 2.26-2.45 (m, 2H), 2.85 (t, J = 8.1 Hz, 2H), 3.97 (t, J = 7.2 Hz, 2H), 5.59 (s, 2H), 6.88 (s, 1H), 7.70-7.89 (m, 2H), 8.04 (dd, J = 8.6 Hz, 1.4 Hz, 1H), 8.60 (s, 1H), 11.77 (s, 1H). ^13^C NMR (500 MHz, DMF-d7, δ): 17.87, 32.82, 45.05, 64.50, 103.02, 111.61, 120.70, 122.60, 123.51, 127.52, 127.98, 139.46, 166.96, 168.32, 176.78. MS m/z 309.2 [M + Na]^+^. HRMS (m/z): [M + Na]^+^ calcd for C_15_H_14_N_2_NaO_4_, 309.0846; found 309.0837.

**Fig. S2e.**
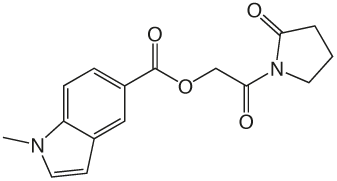

**1l**2-oxo-2-(2-oxopyrrolidin-1-yl)ethyl 1-methyl-1H-indole-5-carboxylate

### 1l

was prepared according to the procedure in General Method A from **1v** and 1-methyl-1H-indole-5-carboxylic acid on a 0.39 mmol scale. The crude product was purified three times via silica gel chromatography using a gradient from 0 to 3% MeOH in CH_2_Cl_2_ to give **1l** (6.3 mg, 0.021 mmol). ^1^H NMR (500 MHz, CDCl_3_, δ): 2.10-2.26 (m, 2H), 2.67 (t, J = 8.1 Hz, 2H), 3.86 (s, 3H), 3.89 (t, J = 7.2 Hz, 2H), 5.44 (s, 2H), 6.62 (d, J = 3.0 Hz, 1H), 7.14 (d, J = 3.2 Hz, 1H), 7.37 (d, J = 8.7 Hz, 1H), 8.02 (dd, J = 8.7 Hz, 1.4 Hz, 1H), 8.51 (s, 1H). ^13^C NMR (500 MHz, CDCl_3_, δ): 17.93, 33.02, 33.07, 44.91, 64.64, 102.78, 108.90, 120.53, 123.31, 124.51, 128.00, 130.26, 139.36, 167.25, 168.55, 176.04. MS m/z 323.2 [M + Na]^+^. HRMS (m/z): [M + Na + CH_3_OH]+1 calcd for C_17_H_20_N_2_NaO_5_, 355.1264; found 355.1258.

**Fig. S2f.**
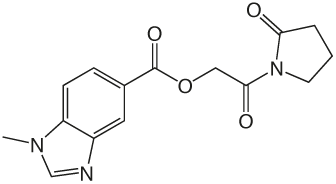

**1m** 2-oxo-2-(2-oxopyrrolidin-1-yl)ethyl 1-methyl-1H-benzo[d]imidazole-5-carboxylate

### 1m

was prepared according to the procedure in General Method A from **1v** and 1-methyl-1H-benzimidazole-5-carboxylic acid on a 0.36 mmol scale under argon. The crude product was purified twice via silica gel chromatography using a gradient from 0 to 15% MeOH in CH_2_Cl_2_ to give **1m** (4 mg, 0.013 mmol). ^1^H NMR (500 MHz, CDCl_3_, δ): 2.11-2.28 (m, 2H), 2.68 (t, J = 8.1 Hz, 2H), 3.81-4.03 (m, 5H), 5.47 (s, 2H), 7.46 (d, J = 8.5 Hz, 1H), 7.99 (s, 1H), 8.15 (dd, J = 8.5 Hz, 1.3 Hz, 1H), 8.64 (s, 1H). ^13^C NMR (500 MHz, CDCl_3_, d): 17.94, 31.28, 33.00, 44.91, 64.88, 109.15, 123.35, 123.65, 124.93, 138.01, 143.40, 145.29, 166.64, 168.25, 176.05. MS m/z 302.2 [M + H]^+^. HRMS (m/z): [M + H]^+^ calcd for C_15_H_16_N_3_O_4_, 302.1135; found 302.1142.

**Fig. S2g.**
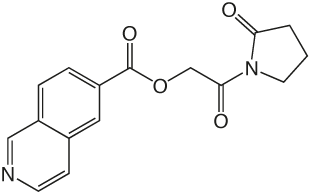

**1n** 2-oxo-2-(2-oxopyrrolidin-1-yl)ethyl isoquinoline-6-carboxylate

### 1n

was prepared according to the procedure in General Method A from **1v** and isoquinoline-6-carboxylic acid on a 0.36 mmol scale. The crude product was purified four times via silica gel chromatography using a gradient from 0 to 10% MeOH in CH_2_Cl_2_ to give **1n** (6.4 mg, 0.021 mmol). ^1^H NMR (500 MHz, CDCl_3_, δ): 2.13-2.31 (m, 2H), 2.69 (t, J = 8.1 Hz, 2H), 3.90 (t, J = 7.1 Hz, 2H), 5.52 (s, 2H), 7.80 (d, J = 5.3 Hz, 1H), 8.09 (d, J = 8.5 Hz, 1H), 8.27 (d, J = 8.4 Hz, 1H), 8.65 (d, J = 5.3 Hz, 1H), 8.70 (s, 1H), 9.39 (s, 1H). ^13^C NMR (500 MHz, CDCl_3_, δ): 17.95, 32.94, 44.91, 65.27, 121.45, 126.98, 128.04, 129.96, 130.90, 135.02, 143.91, 152.57, 165.55, 167.74, 176.15. MS m/z 321.3 [M + Na]^+^. HRMS (m/z): [M + H]^+^ calcd for C_16_H_15_N_2_O_4_, 299.1026; found 299.1027.

**Fig. S2h.**
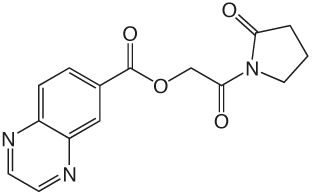

**1o** 2-oxo-2-(2-oxopyrrolidin-1-yl)ethyl quinoxaline-6-carboxylate **8a.** was prepared according to the procedure in General Method A from **1v** and quinoxaline-6-carboxylic acid on a 0.37 mmol scale. The crude product was purified via silica gel chromatography using a gradient from 0 to 10% MeOH in CH_2_Cl_2_ to give **1o** (20.6 mg, 0.069 mmol). ^1^H NMR (500 MHz, CDCl_3_, δ): 2.08-2.29 (m, 2H), 2.67 (t, J = 7.6 Hz, 2H), 3.88 (t, J = 6.2 Hz, 2H), 5.51 (s, 2H), 8.20 (d, J = 8.4 Hz, 1H), 8.44 (d, J = 8.1 Hz, 1H), 8.86-9.06 (m, 3H). ^13^C NMR (500 MHz, CDCl_3_, δ): 17.94, 32.92, 44.90, 65.35, 129.75, 129.93, 130.77, 132.80, 142.30, 145.15, 146.01, 146.75, 165.24, 167.64, 176.14. MS m/z 322.3 [M + Na]^+^. HRMS (m/z): [M + Na]^+^ calcd for C_15_H_13_N_3_NaO_4_, 322.0798; found 322.0790.

**Fig. S2i.**
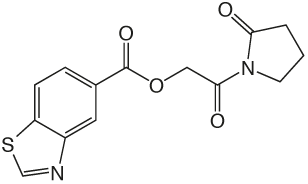

**1p** 2-oxo-2-(2-oxopyrrolidin-1-yl)ethyl benzothiazole-5-carboxylate

### 1p

was prepared according to the procedure in General Method A from **1v** and 5-benzothiazole carboxylic acid on a 0.36 mmol scale. The crude product was purified via silica gel chromatography using a gradient from 0 to 3% MeOH in CH_2_Cl_2_ to give **1p** (43.7 mg, 0.14 mmol). ^1^H NMR (500 MHz, CDCl_3_, δ): 2.06-2.25 (m, 2H), 2.64 (t, J = 8.1 Hz, 2H), 3.85 (t, J = 7.2 Hz, 2H), 5.46 (s, 2H), 8.03 (d, J = 8.4 Hz, 1H), 8.18 (d, J = 7.6 Hz, 1H), 8.88 (s, 1H), 9.09 (s, 1H). ^13^C NMR (500 MHz, CDCl_3_, δ): 17.91, 32.93, 44.89, 65.11, 121.93, 125.64, 126.36, 127.86, 138.91, 153.13, 155.39, 165.85, 167.86, 176.13. MS m/z 327.3 [M + Na]^+^. HRMS (m/z): [M + Na]^+^ calcd for C_14_H12N2NaO_4_S, 327.0410; found 327.0407.

**Fig. S3.**
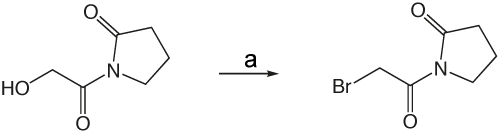

**Scheme 3. Synthesis of 1x.** a).CBr4, triphenylphospine, CH_2_Cl_2_, r.t. overnight.

**Fig. S3a.**
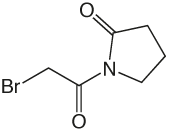

**1x** 1-(2-Bromoacetyl)pyrrolidin-2-one

### 1x

Carbon tetrabromide (126 mg, 0.38 mmol) was added to a solution of **1v** (50 mg, 0.35 mmol) in 1 mL CH_2_Cl_2_ at 0 °C. A solution of triphenylphosphine (100 mg, 0.38 mmol) in 0.5 mL CH_2_Cl_2_ was added dropwise and the reaction mixture allowed to warm to room temperature overnight. The crude reaction mixture was concentrated in vacuo and purified via silica gel chromatography using a gradient from 0 to 5% MeOH in CH_2_Cl_2_ to give **1x** (48 mg, 0.23 mmol). ^1^H NMR (500 MHz, CDCl_3_, δ): 2.05-2.17 (m, 2H), 2.65 (t, J = 8.1 Hz, 2H), 3.86 (t, J = 7.3 Hz, 2H), 4.50 (s, 2H). ^13^C NMR (500 MHz, CDCl_3_, δ): 17.35, 30.09, 33.11, 45.82, 166.61, 175.38. MS m/z 230.2 [M + Na]^+^. HRMS (m/z): [M + Na + CH_3_OH]+1 calcd for C_7_H_12_BrNNaO_3_, 259.9893; found 259.9892.

### Scheme 4. Synthesis of 1s. a)

**Fig. S4.**
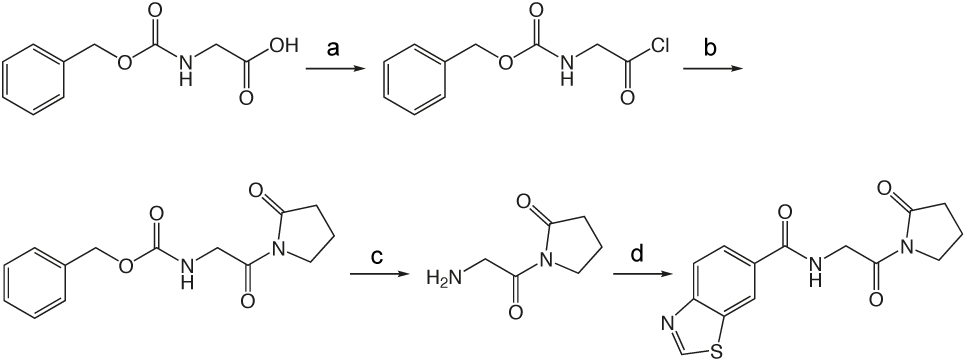

PCl_5_, ether, −15C - 0C, 3h. b). TMS-2-pyrrolidinone, ether, 0°C-r.t. 1.25h. c). TFA, thioanisole, TMSBr, r.t. 1.5h. d). benzohiazole-6-carboxylic acid, HBTU, HOAt, DIEA, DMF, CH_2_Cl_2_, r.t. overnight.

**Fig. S4a.**
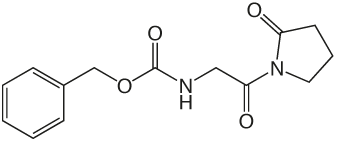

**1y** N-(2-Oxo-2-(2-oxopyrrolidin-1-yl)ethyl)benzo[d]thiazole-6-carboxamide.

### 1y

To a solution of Cbz-Gly (418 mg, 2.0 mmol) in 2.5 mL ether at −15 °C was added PCl5 (458 mg, 2.2 mmol). The reaction mixture was stirred for 3 h at 0 °C, and then concentrated in vacuo. The crude acid chloride was taken up in 3.5 mL ether, and 1-TMS-2-pyrrolidinone (0.35 mL, 2.2 mmol) in 3.5 mL ether was added dropwise at 0 °C. The ice bath was removed and the reaction mixture stirred at room temperature 1.25 h, then concentrated in vacuo. The crude product was partitioned between ethyl acetate and 10% NaHCO_3_ solution. The aqueous was reextracted three times with ethyl acetate, and the combined organic extracts washed with brine, dried over Na2SO4, filtered and concentrated in vacuo. The crude product was purified via silica gel chromatography using a gradient from 0 to 10% MeOH in CH_2_Cl_2_ to give **1y** (369 mg, 1.34 mmol). ^1^H NMR (500 MHz, CDCl_3_, δ): 1.85-2.07 (m, 2H), 2.47 (t, J = 8.1 Hz, 2H), 3.68 (t, J = 7.2 Hz, 2H), 4.42 (d, J = 5.6 Hz, 2H), 5.05 (s, 2H), 5.88 (br t, J = 5.2 Hz, 1H), 7.15-7.50 (m, 5H). ^13^C NMR (500 MHz, CDCl_3_, δ): 17.52, 33.01, 45.10, 46.33, 66.72, 127.96, 128.29, 136.56, 156.56, 170.39, 175.85. MS m/z 299.3 [M + Na]^+^.

**Fig. S4b.**
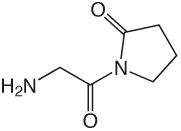

**1z** 1-(2-aminoacetyl)pyrrolidin-2-one

### 1z

To a solution of **1y** (187 mg, 0.68 mmol) in 13.5 mL TFA was added thioanisole (4.0 mL, 33.88 mmol) and TMSBr (0.89 mL, 6.78 mmol). The reaction mixture stirred at room temperature for 1.5 h, then concentrated in vacuo. The crude product was purified via silica gel chromatography using a gradient from 0 to 25% MeOH in CH_2_Cl_2_ to give **1z** which was used without further purification. ^1^H NMR (500 MHz, MeOD, δ): 2.08-2.25 (m, 2H), 2.33 (t, J = 8.1 Hz, 1H), 2.66 (t, J = 8.1 Hz, 1H), 3.43 (t, J = 7.0 Hz, 1H), 3.77-3.94 (m, 3H), 4.35 (br s, 2H). ^13^C NMR (500 MHz, MeOD, δ): 17.23, 32.31, 42.30, 44.88, 167.37, 177.17. MS m/z 143.2 [M + H]^+^.

**Fig. S4c.**
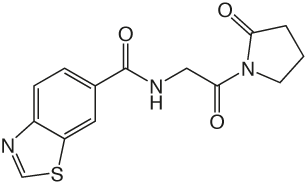

**1s** N-(2-oxo-2-(2-oxopyrrolidin-1-yl)ethyl)-benzo[d]thiazole-6-carboxamide

### 1s

A solution of benzothiazole-6-carboxylic acid (122 mg, 0.68 mmol), HBTU (774 mg, 2.04 mmol), and HOAt (278 mg, 2.04 mmol) in 5 mL CH_2_Cl_2_ and 1 mL DMF was stirred at room temperature 1 h, then **1z** (96 mg, 0.68 mmol) and DIEA (0.53 mL, 3.04 mmol) in 2 mL DMF was added and the reaction mixture stirred overnight. The reaction mixture was diluted with CH_2_Cl_2_ and washed with water and saturated NaHCO_3_ solution, dried over Na2SO4, filtered and concentrated in vacuo. The crude product was purified via reverse-phase HPLC using a gradient from 10 to 95% B in A over 30 min to give **1s** (20.5 mg, 0.068 mmol). ^1^H NMR (500 MHz, CDCl_3_, d): 2.10-2.25 (m, 2H), 2.67 (t, J = 8.1 Hz, 2H), 3.89 (t, J = 7.2 Hz, 2H), 4.84 (d, J = 5.1 Hz, 2H), 7.25 (br s, 1H), 7.97 (dd, J = 8.5 Hz, 1.6 Hz, 1H), 8.18 (d, J = 8.5 Hz, 1H), 8.53 (d, J = 1.4 Hz, 1H), 9.18 (s, 1H). ^13^C NMR (500 MHz, CDCl_3_, δ): 17.67, 33.09, 45.21, 45.67, 121.99, 123.42, 124.98, 131.42, 134.03, 154.79, 157.08, 166.87, 170.01, 175.71. MS m/z 304.4 [M + H]^+^. HRMS (m/z): [M + H]^+^ calcd for C_14_H_14_N_3_O_3_S, 304.0750; found 304.0751.

### Scheme 5. Synthesis of 1t

**Fig. S5.**
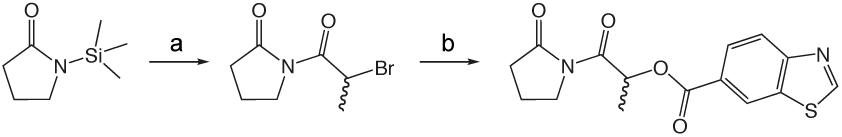

a). 2-bromopropionyl bromide, ether 0°C - r.t. 1.25h. b). benzothiazole-6-carboxylic acid, Cs_2_CO_3_, DMF r.t. overnight.

**Fig. S5a.**
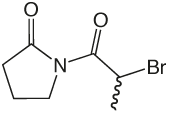

**1aa** 1-(2-bromopropanoyl)pyrrolidin-2-one.

### 1aa

To a solution of 2-bromopropionyl bromide (100 µL, 0.95 mmol) in 2 mL ether at 0 °C was added dropwise quickly a solution of 1-TMS-2-pyrrolidinone (168 µL, 1.05 mmol) in 2 mL ether. The ice bath was removed and the reaction mixture stirred at room temperature 1.25 h, then concentrated in vacuo. The residue was partitioned between ethyl acetate and 10% NaHCO_3_. The aqueous layer was reextracted three times with ethyl acetate and the combined organic layers washed with brine, dried over Na2SO4, filtered, and concentrated in vacuo to give **1aa** (191.7 mg, 0.87 mmol) which was used without further purification. ^1^H NMR (500 MHz, CDCl_3_, δ): 1.76 (d, J = 6.8 Hz, 3H), 1.99-2.10 (m, 2H), 2.53-2.67 (m, 2H), 3.71-3.89 (m, 2H), 5.63 (q, J = 6.8 Hz, 1H). ^13^C NMR (500 MHz, CDCl_3_, δ): 17.05, 20.62, 33.51, 40.44, 46.03, 170.29, 174.92. MS m/z 244.0 [M + Na]^+^.

**Fig. S5b.**
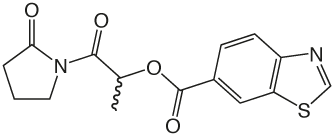

**1t** 1-oxo-1-(2-oxopyrrolidin-1-yl)propan-2-yl benzo[d]thiazole-6-carboxylate.

### 1t

To a solution of **1aa** (192 mg, 0.87 mmol) in 4 mL DMF was added benzothiazole-6-carboxylic acid (156 mg, 0.87 mmol) and cesium carbonate (311 mg, 0.95 mmol). The reaction mixture stirred overnight to completion and was partitioned between H_2_O and CH_2_Cl_2_. The aqueous layer was reextracted twice with CH_2_Cl_2_ and the combined organic layers washed with brine, dried over Na2SO4, filtered and concentrated in vacuo. The crude product was purified via silica gel chromatography using a gradient from 40 to 60% EtOAc in hexanes to give **1t** (184 mg, 0.58 mmol). ^1^H NMR (500 MHz, CDCl_3_, d): 1.64 (d, J = 6.8 Hz, 3H), 1.96-2.17 (m, 2H), 2.48-2.77 (m, 2H), 3.67-3.98 (m, 2H), 6.20 (q, J =6.8 Hz, 1H), 8.09-8.27 (m, 2H), 8.71 (d, J = 1.1 Hz, 1H), 9.14 (s, 1H). ^13^C NMR (500 MHz, CDCl_3_, δ): 16.63, 17.48, 33.35, 45.45, 70.89, 123.39, 124.52, 126.78, 127.50, 133.74, 156.20, 157.61, 165.56, 171.70, 175.30. MS m/z 341.2 [M + Na]^+^. HRMS (m/z): [M + H]^+^ calcd for C_15_H_15_N_2_O_4_S, 319.0747; found 319.0743.

**Fig. S6.**
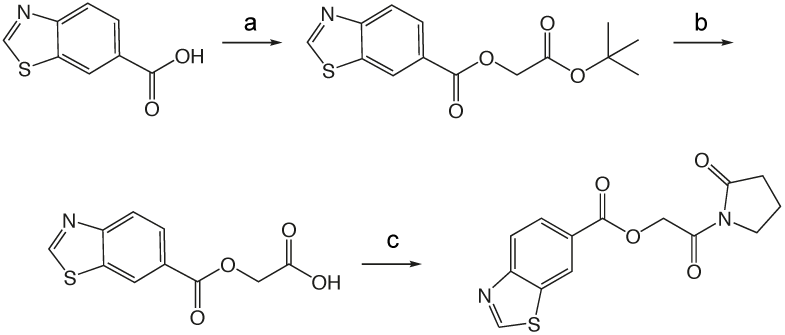

**Scheme 6. Synthesis of 1u.** a). t-butyl 3-hydroxypropionate, EDCI, DMAP, CH_2_Cl_2_, DMF, r.t. overnight. b). Pd-C, EtOH, DMF, H_2_ c). biphenyl-4-carboxylic acid, EDCI, DMAP, CH_2_Cl_2_, DMF, r.t. overnight.

**Fig. S6a.**
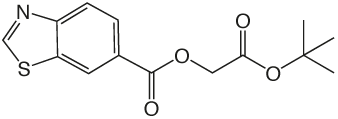

**1ab.** 3-tert-butoxy-3-oxopropylbenzo[d]thiazole-6-carboxylate

### 1ab

was prepared according to the procedure in General Method A from tert-butyl 3-hydroxypropionate and benzothiazole-6-carboxylic acid on a 1.0 mmol scale. The crude product was purified via silica gel chromatography using a gradient from 0 to 5% MeOH in CH_2_Cl_2_ to give **1ab** (227 mg, 0.74 mmol). ^1^H NMR (500 MHz, CDCl_3_, δ): 1.40 (s, 9H), 2.69 (t, J = 6.3 Hz, 2H), 4.56 (t, J = 6.3 Hz, 2H), 8.09 (s, 2H), 8.60 (s, 1H), 9.11 (s, 1H). ^13^C NMR (500 MHz, CDCl_3_, δ): 28.02, 35.25, 61.07, 81.04, 123.33, 124.21, 127.20, 127.26, 133.73, 156.02, 157.43, 165.63, 169.80. MS m/z 330.3 [M + Na]^+^.

**Fig. S6b.**
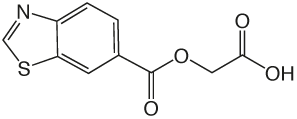

**1ac.** 3-(benzo[d]thiazole-6-carbonyloxy)propanoic acid

### 1ac

Compound **1ab** (227 mg, 0.74 mmol) was dissolved in 3 mL TFA and 3 mL CH_2_Cl_2_, stirred at room temperature for 2.5 h and concentrated in vacuo to give **1ac** (171 mg, 0.68 mmol), which was used without further purification. ^1^H NMR (500 MHz, CDCl_3_, δ): 2.94 (t, J = 6.0 Hz, 2H), 4.70 (t, J = 6.0 Hz, 2H), 8.21 (s, 2H), 8.73 (s, 1H), 9.22 (s, 1H). MS m/z 252.3 [M + H]^+^.

**Fig. S6c.**
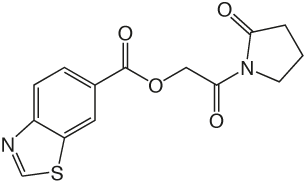

**1u.** 3-oxo-3-(2-oxopyrrolidin-1-yl)propyl benzo[d]thiazole-6-carboxylate

### 1u

Compound **1ac** (170 mg, 0.68 mmol) was taken up in 5 mL thionyl chloride with a drop of DMF and stirred at room temperature overnight, then concentrated in vacuo. The crude acid chloride was taken up in 7 mL THF and chilled to 0 °C in an ice bath. 1-TMS-2-pyrrolidinone (0.12 mL, 0.75 mmol) in 1 mL THF was added dropwise quickly, the ice bath removed, and the reaction mixture stirred at room temperature 4 h before concentrating in vacuo. The crude product was purified via silica gel chromatography using a gradient from 0 to 5% MeOH in CH_2_Cl_2_ to give **1u** (108 mg, 0.34 mmol). ^1^H NMR (500 MHz, CDCl_3_, δ): 1.95-2.13 (m, 2H), 2.59 (t, J = 8.0 Hz, 2H), 3.40 (t, J = 6.2 Hz, 2H), 3.81 (t, J = 7.1 Hz, 2H), 4.69 (t, J = 6.2 Hz, 2H), 8.04-8.22 (m, 2H), 8.63 (s, 1H), 9.13 (s, 1H). ^13^C NMR (500 MHz, CDCl_3_, δ): 17.20, 33.50, 36.38, 45.34, 60.21, 123.31, 124.27, 127.32, 127.38, 133.70, 156.01, 157.42, 165.79, 170.91, 175.62. MS m/z 341.3 [M + Na]^+^. HRMS (m/z): [M + Na]^+^ calcd for C_15_H_14_N_2_NaO_4_S, 341.0566; found 341.0562.

**Fig. S6d.**
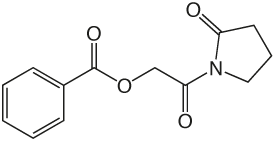

**1h.** 2-oxo-2-(2-oxopyrrolidin-1-yl)ethyl benzoate.

### 1h

was prepared according to the procedure in General Method A from **1v** and benzoic acid on a 0.35 mmol scale. The crude product was purified via silica gel chromatography using a gradient from 0 to 3% MeOH in CH_2_Cl_2_ to give **1h** (52.1 mg, 0.21 mmol). ^1^H NMR (500 MHz, CDCl_3_, δ): 2.02-2.18 (m, 2H), 2.61 (t, J = 8.1 Hz, 2H), 3.83 (t, J = 7.2 Hz, 2H), 5.40 (s, 2H), 7.46 (t, J = 7.7 Hz, 2H), 7.58 (t, J = 7.4 Hz, 1H), 8.11 (d, J = 8.3 Hz, 2H). ^13^C NMR (500 MHz, CDCl_3_, δ): 17.87, 32.90, 44.86, 64.82, 128.40, 129.48, 129.92, 133.28, 166.11, 167.96, 176.05. MS m/z 270.3 [M + Na]^+^. HRMS (m/z): [M + Na]^+^ calcd for C_13_H_13_NNaO_4_, 270.0742; found 270.0741.

**Fig. S6e.**
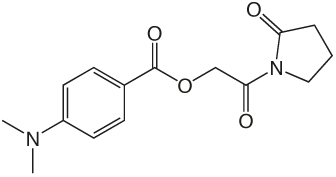

**1g.** 2-oxo-2-(2-oxopyrrolidin-1-yl)ethyl 4-(dimethylamino)benzoate

### 1g

was prepared according to the procedure in General Method A from **1v** and 4-(dimethylamino)-benzoic acid on a 0.35 mmol scale. The crude product was purified via silica gel chromatography using a gradient from 0 to 10% MeOH in CH_2_Cl_2_ to give **1q** (3.1 mg, 0.011 mmol). ^1^H NMR (500 MHz, CDCl_3_, δ): 2.09-2.23 (m, 2H), 2.65 (t, J = 8.1 Hz, 2H), 3.07 (s, 6H), 3.87 (t, J = 7.2 Hz, 2H), 5.36 (s, 2H), 6.69 (d, J = 9.0 Hz, 2H), 8.00 (d, J = 9.0 Hz, 2H). ^13^C NMR (500 MHz, CDCl_3_, δ): 17.90, 32.99, 40.12, 44.88, 64.32, 110.69, 116.05, 131.74, 153.56, 166.39, 168.68, 175.99. MS m/z 291.1 [M + H]^+^. HRMS (m/z): [M + H]^+^ calcd for C_15_H_19_N_2_O_4_, 291.1345; found 291.1348

**Fig. S6f.**
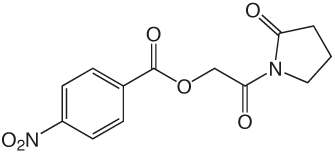

**1r.** 2-oxo-2-(2-oxopyrrolidin-1-yl)ethyl 4-nitrobenzoate

### 1r

was prepared according to the procedure in General Method A from **1v** and 4-nitrobenzoic acid on a 0.35 mmol scale. The crude product was purified via silica gel chromatography using a gradient from 0 to 3% MeOH in CH_2_Cl_2_ to give **1r** (50.9 mg, 0.17 mmol). ^1^H NMR (500 MHz, CDCl_3_, δ): 2.02-2.25 (m, 2H), 2.64 (t, J = 8.1 Hz, 2H), 3.84 (t, J = 7.2 Hz, 2H), 5.43 (s, 2H), 8.05-8.36 (m, 4H). ^13^C NMR (500 MHz, CDCl_3_, δ): 17.91, 32.85, 44.87, 65.40, 123.57, 131.06, 134.85, 150.72, 164.29, 167.34, 176.20. MS m/z 315.2 [M + Na]^+^. HRMS (m/z): [M + Na]^+^ calcd for C_13_H_12_O_6_N_2_Na, 315.0588; found 315.0594.

**Fig. S6g.**
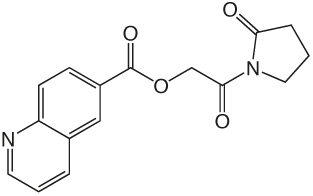

**1ad.** 2-oxo-2-(2-oxopyrrolidin-1-yl)ethyl quinoline-6-carboxylate

### 1ad

was prepared according to the procedure in General Method A from **1v** and quinolone 6-carboxylic acid on a 0.36 mmol scale. The crude product was purified via silica gel chromatography using a gradient from 0 to 10% MeOH in CH_2_Cl_2_ to give **1ad** (26.6 mg, 0.089 mmol). ^1^H NMR (500 MHz, CDCl_3_, δ): 2.09-2.32 (m, 2H), 2.66 (t, J = 8.1 Hz, 2H), 3.88 (t, J = 7.2 Hz, 2H), 5.49 (s, 2H), 7.42-7.63 (m, 1H), 8.18 (d, J = 8.8 Hz, 1H), 8.29 (d, J = 7.6 Hz, 1H), 8.38 (dd, J = 8.8 Hz, 1.8 Hz, 1H), 8.70 (d, J = 1.4 Hz, 1H), 8.97-9.16 (m, 1H). ^13^C NMR (500 MHz, CDCl_3_, δ): 17.91, 32.91, 44.88, 65.10, 121.87, 127.42, 127.50, 129.13, 129.90, 131.52, 137.38, 150.28, 152.63, 165.64, 167.85, 176.04. MS m/z 299.2 [M + H]^+^. HRMS (m/z): [M + Na]^+^ calcd for C_16_H_14_N_2_NaO_4_, 321.0846; found 321.0846.

**Fig. S6h.**
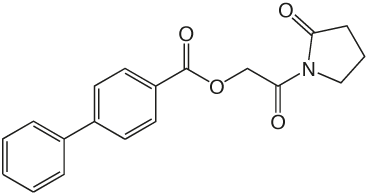

**1ae.**2-oxo-2-(2-oxopyrrolidin-1-yl)ethyl biphenyl-4-carboxylate

### 1ae

was prepared according to the procedure in General Method A from **1v** and biphenyl-4-carboxylic acid on a 0.36 mmol scale. The crude product was purified via silica gel chromatography using a gradient from 0 to 3% MeOH in CH_2_Cl_2_ to give **1ae** (49.2 mg, 0.15 mmol). ^1^H NMR (500 MHz, CDCl_3_, δ): 1.82-1.99 (m, 2H), 2.40 (t, J = 8.1 Hz, 2H), 3.62 (t, J = 7.2 Hz, 2H), 5.07 (s, 2H), 7.17 (t, J = 7.3 Hz,1H), 7.25 (t, J = 7.5 Hz, 2H), 7.41 (d, J = 7.4 Hz, 2H), 7.46 (d, J = 8.3 Hz, 2H), 7.96 (d, J = 8.3 Hz, 2H). ^13^C NMR (500 MHz, CDCl_3_, δ): 18.13, 33.17, 45.11, 65.10, 127.32, 127.54, 128.44, 129.18, 130.72, 140.18, 146.21, 166.23, 168.23, 176.33. MS m/z 346.3 [M + Na]^+^. HRMS (m/z): [M + Na]^+^ calcd for C_19_H_17_NNaO_4_, 346.1050; found 346.1052.

## Supplementary 2: Tables

**Supplementary Table 1.**
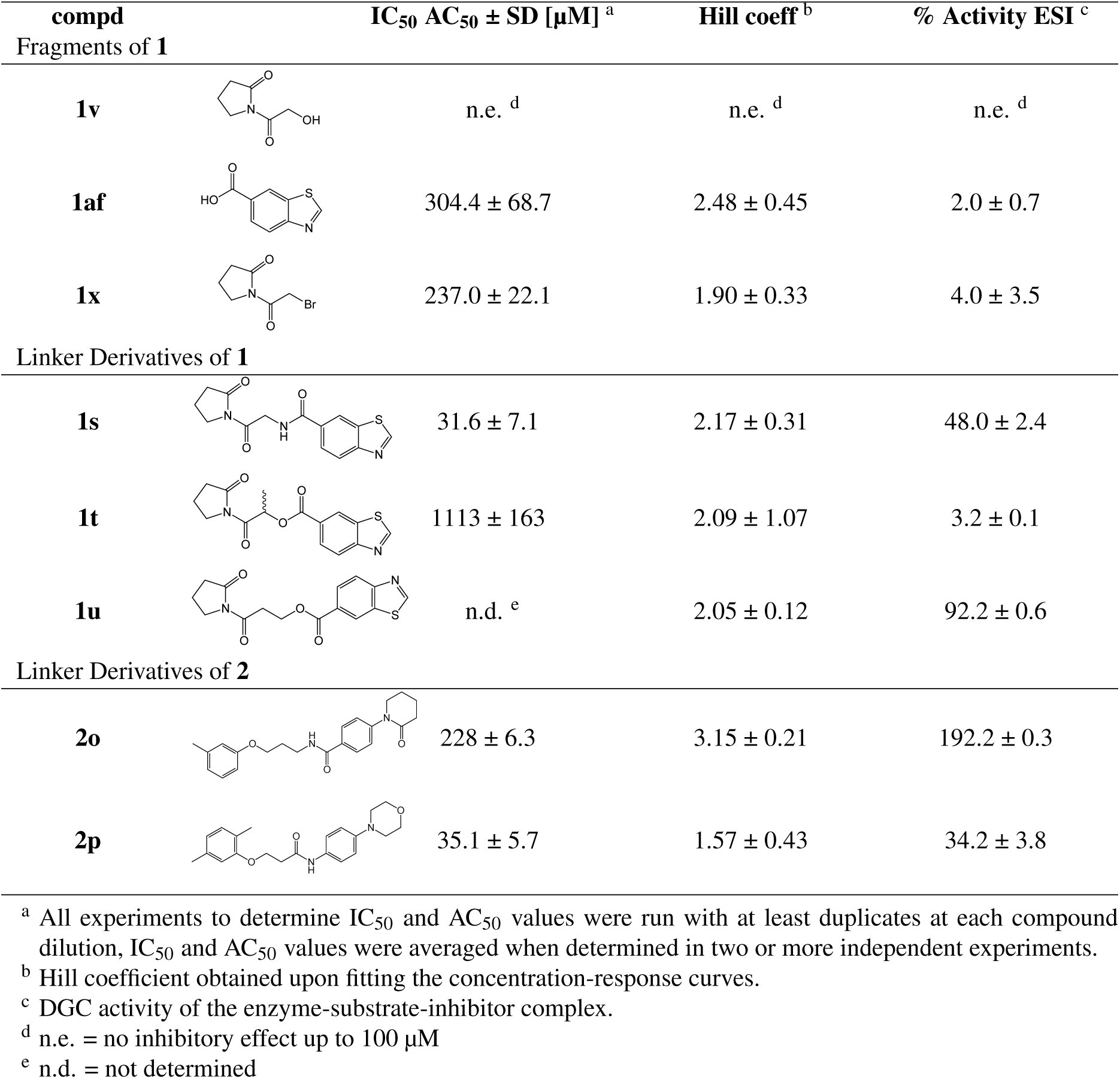
Fragment analysis of and linker substituents of **1** and **2**.

| compd |  | IC<sub>50</sub> AC<sub>50</sub> ± SD [μM]<sup>a</sup> | Hill coeff<sup>b</sup> | % Activity ESI<sup>c</sup> |
| --- | --- | --- | --- | --- |
| Fragments of **1** |  |  |  |  |
| **1v** |  | n.e.<sup>d</sup> | n.e.<sup>d</sup> | n.e.<sup>d</sup> |
| **1af** |  | 304.4 ± 68.7 | 2.48 ± 0.45 | 2.0 ± 0.7 |
| **1x** |  | 237.0 ± 22.1 | 1.90 ± 0.33 | 4.0 ± 3.5 |
| Linker Derivatives of **1** |  |  |  |  |
| **1s** |  | 31.6 ± 7.1 | 2.17 ± 0.31 | 48.0 ± 2.4 |
| **1t** |  | 1113 ± 163 | 2.09 ± 1.07 | 3.2 ± 0.1 |
| **1u** |  | n.d.<sup>e</sup> | 2.05 ± 0.12 | 92.2 ± 0.6 |
| Linker Derivatives of **2** |  |  |  |  |
| **2o** |  | 228 ± 6.3 | 3.15 ± 0.21 | 192.2 ± 0.3 |
| **2p** |  | 35.1 ± 5.7 | 1.57 ± 0.43 | 34.2 ± 3.8 |

<sup>a</sup> All experiments to determine IC<sub>50</sub> and AC<sub>50</sub> values were run with at least duplicates at each compound dilution, IC<sub>50</sub> and AC<sub>50</sub> values were averaged when determined in two or more independent experiments.
<sup>b</sup> Hill coefficient obtained upon fitting the concentration-response curves.
<sup>c</sup> DGC activity of the enzyme-substrate-inhibitor complex.
<sup>d</sup> n.e. = no inhibitory effect up to 100 μM
<sup>e</sup> n.d. = not determined

**Supplementary Table 2.**
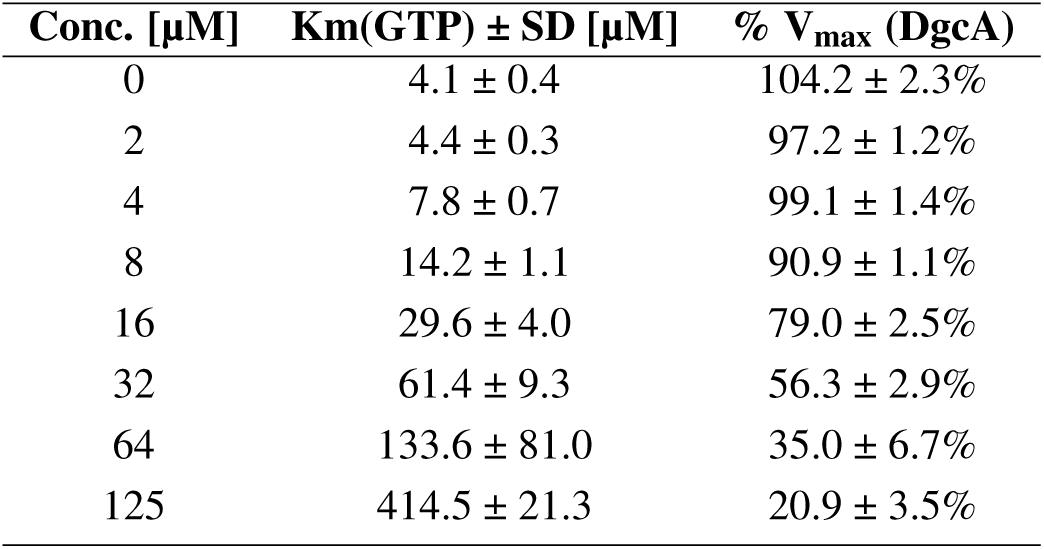
Effect of **1** on DgcA GTP binding and maximal rate of DGC reaction.

**Supplementary Table 3.** Effect of **2** on DgcA GTP binding and maximal rate of DGC reaction.

| Conc. [$\mu\text{M}$] | Km(GTP) ± SD [$\mu\text{M}$] | % V<sub>max</sub> (DgcA) |
| --- | --- | --- |
| 0 | 5.61 ± 1.51 | 96.0 ± 5.3% |
| 2 | 3.86 ± 0.94 | 105.5 ± 5.6% |
| 4 | 5.50 ± 1.53 | 112.1 ± 9.1% |
| 8 | 7.47 ± 2.25 | 106.8 ± 9.0% |
| 16 | 14.46 ± 2.87 | 107.7 ± 3.9% |
| 32 | 56.13 ± 11.47 | 103.4 ± 5.1% |
| 64 | 282.52 ± 11.52 | 119.6 ± 20.7% |
| 125 | 934.63 ± 64.08 | 102.8 ± 28.4% |

